# A Comprehensive Analysis of Atlantic Salmon Gonad and Pituitary Transcriptomes Identifies Novel Players in Sexual Maturation

**DOI:** 10.1101/2025.06.16.659895

**Authors:** Xin-Di Huang, Morgane Frapin, Ehsan Pashay Ahi, Craig Robert Primmer, Jukka-Pekka Verta

## Abstract

**Background:** Sexual maturation is a key developmental process important for reproductive success. Understanding the molecular mechanisms behind variation in sexual maturation can provide insights into reproductive biology and how life history variation is encoded in the genome. Atlantic salmon (*<u>Salmo salar</u>*) has become an excellent sexual maturation research model due to its diversity of life history strategies and its ecological and economic importance. A major challenge has been the lack of a comprehensive transcriptional investigation of reproductive tissues that captures the dynamic transcriptional changes across individuals, tissues, and developmental stages. Long non-coding RNAs (lncRNAs) also play crucial roles in maturation, yet their functions in salmon maturation remain underexplored.

**Results:** In this study, we sequenced 98 transcriptomes and found substantial transcriptomic complexity in the gonad and pituitary tissues of Atlantic salmon. We identified transcripts corresponding to 2,364 putative newly characterized protein-coding genes and 4,421 putative long intergenic non-coding RNAs (lincRNAs), many with tissue-specific expression. Gene co-expression network analysis (WGCNA) revealed tissue-specific gene network modules, linked to GO terms including Wnt signaling in immature testis, lipid metabolism, and cilia assembly in mature testis, ribosome biogenesis and DNA repair in the ovary, and hormone activity in pituitary. We identified new copies of known genes, such as *gh1, pou3f2*, and *ier5* associated with the regulation of gonadal and pituitary functions. Some lncRNAs and their nearest genes showed correlated expression within modules, suggesting potential regulatory roles. Candidate lincRNAs indicated cis-acting regulatory potential on genes like *tnfrsf11b* and *fgl1*, which are implicated in immune privilege during gonadal development and sperm quality control.

**Conclusions:** Our study provides a comprehensive transcriptomic analysis of Atlantic salmon gonad and pituitary tissues, significantly improving the functional annotation of the Atlantic salmon genome. These findings reveal key regulatory pathways and novel molecular players involved in sexual maturation, particularly in the testis. Importantly, our study highlights the regulatory potential of lncRNAs in reproductive biology and maturation age variation, advancing our understanding of the molecular mechanisms governing sexual maturation. They further unlock future gene expression analyses and regulatory network reconstruction for dissecting the roles of lncRNAs in Atlantic salmon life history variation.

## Background

The age at which an individual reaches sexual maturity is a polymorphic life history trait with significant fitness consequences [1, 2]. Variation in maturation age contributes to variation in the life history strategies both within and among species [3, 4]. Delayed maturity provides more opportunities to grow larger body size and accumulate higher energy reserves for reproduction, potentially increasing fecundity and offspring survival, however, it also increases the risk of mortality before reproduction and extends the generation time [5]. Following from their indeterminate growth, larger, older fish individuals contribute disproportionately to reproduction, making maturation age variation especially important in determining population growth, resilience, and evolutionary responses in fishes [6]. Thus, a better understanding of the mechanisms that encode for variation in maturation age not only gives insight into the evolutionary trade-off between reproduction and survival, but also holds significant potential for practical applications, such as in conservation and aquaculture.

Atlantic salmon (*Salmo salar*, Salmonidae) has become an excellent research model for age-at-maturity because of its life history diversity and its ecological and economic significance [1]. Atlantic salmon exhibits incredibly high diversity in life history strategies, for example, 120 life history strategies were reported over a 40-year period in a single river system [7]. As an important farmed seafood species, early puberty of Atlantic salmon can lead to slower growth and susceptibility to disease, which is detrimental to fish health and product quality [8–10]. Thus, a deeper understanding of the regulatory mechanisms controlling sexual maturation in Atlantic salmon is valuable for improving breeding and aquaculture practices.

As in other vertebrates, maturation in Atlantic salmon is a complex process regulated by the brain-pituitary-gonadal (BPG) axis [1, 11]. This axis orchestrates hormonal control, gene expression, and gonadal development, making it a crucial focus for understanding the mechanisms of sexual maturation regulation [10]. An improved understanding of this process could be gained with an integrative approach that combines transcriptomic data from multiple BPG axis tissues, as transcriptomes provide insight into the molecular mechanisms underlying maturation, with a particular focus on the regulatory roles of individual tissues of the BPG axis [12, 13].

The ENCODE project revealed that while 80% of the human genome is transcribed, less than 2% encodes proteins, with the remaining transcripts classified as non-coding RNAs (ncRNAs), including microRNAs (miRNAs) and long non-coding RNAs (lncRNAs) [14, 15]. Although lncRNAs are biochemically similar to messenger RNAs (mRNAs), they generally do not encode for protein products [16]. LncRNAs tend to have lower expression levels that are more specific to certain tissues, and they show greater variation in expression between individuals, compared to mRNAs [17]. In recent years, many studies on a variety of organisms have shown that non-coding RNAs directly or indirectly regulate the reproductive system [18]. In Atlantic salmon, non-coding RNAs such as miRNAs have been implicated in testis maturation [19]. While the role of lncRNAs in sexual maturation remains less well understood, previous studies have provided valuable insights by linking lncRNAs to smoltification [20] and immunomodulation [21].

One of the major challenges in studying transcriptional regulation using RNA-seq is the gap between known and unknown variation in transcriptomes. Reference genomes often fail to capture the full diversity of transcripts that can arise due to genetic, transcriptional, and post-transcriptional variation [22]. This is particularly true for the testis transcriptome, which is highly complex and serves as a fertile ground for the discovery of novel genes [23, 24]. Earlier transcriptomic studies of Atlantic salmon gondads have predominantly utilized the reference gene model to investigate gene expression [13, 25–27]. While this approach has advanced our understanding of key gene functions, it may offer limited opportunities for the identification of novel genes and lncRNAs. The present study builds on these efforts by exploring beyond the reference annotation to uncover additional regulatory elements involved in sexual maturation. Therefore, align-then-assemble strategies are useful for uncovering hidden transcript diversity [28], especially in studies involving the BPG axis, before variation between individuals can be quantified.

In this study, we applied a reference-based transcriptome assembly to generate a comprehensive view of Atlantic salmon reproductive transcriptomes. This work represents a large-scale analysis of newly characterized protein-coding genes (mostly copies of annotated genes) and lncRNAs in key reproductive tissues of Atlantic salmon, including the testis, ovary, and pituitary. By examining gene expression patterns across these tissues, we aim to provide fundamental insights into the molecular regulation of sexual maturation. Our study not only expands the current understanding of the genetic mechanisms underlying maturation in Atlantic salmon but also identifies newly characterized protein-coding genes and lncRNAs that may play crucial regulatory roles in this process. These findings offer valuable implications for both basic research into reproductive biology and practical applications in aquaculture.

## Materials and methods

### Sample materials

The broodstock is a first-generation hatchery stock composed of a mixture of northern Baltic Sea lineages and maintained by the Natural Resources Institute Finland (LUKE). Eggs were fertilized and raised in controlled conditions at the University of Helsinki research animal facility. Fish rearing and genotyping were described in detail in previous studies [29–31]. In brief, we used Atlantic salmon from two age cohorts that were raised in common garden hatchery conditions, with an annual cycle of natural water temperature and light corresponding to the location in southern Finland. The fish represent a mix of Baltic Sea lineages from Finnish rivers. 1-year-old individuals used in this study were raised in the fish facilities at the Viikki campus of the University of Helsinki, and the 4-year-old individuals used were reared in larger experimental facilities at Lammi Biological Station (University of Helsinki). All experiments were conducted under animal experiment permits granted by the Finnish Project Authorization Board (permits ESAVI/2778/2018 and ESAVI/4511/2020).

Fish were euthanized by using an overdose of buffered tricaine methanesulphonate (MS-222) and dissected. Maturation status was determined based on the presence or absence of milt release. Tissue samples including pituitary and gonads were flash-frozen in liquid nitrogen and stored at -80°C until RNA extraction. Eight pituitary samples were collected from 4-year-old immature males and 90 gonadal samples were collected from 1-year-old individuals, including immature ovaries (n=36), immature testis (n=28), and mature testis (n=26).

### RNA extraction and quantification

Before RNA extraction, mature testis tissues were stripped of most sperm by thawing them at room-temperature in phosphate-buffered saline (PBS) for around 1 minute then using a micro spatula to squeeze while rolling the tissue on a filter paper until only solid tissue remained. Samples were then homogenized in Nucleospin 96 RNA extraction buffer using a Bead Ruptor Elite (Omni Inc, Kennesaw, GA), 2 ml tubes, and 2.4 mm steel beads. Total RNA was extracted from each sample using NucleoSpin 96 RNA kit (MACHEREY-NAGEL, Dueren, Germany). Residual DNA was removed with TURBO DNase (Invitrogen, Waltham, MA). RNA quantity was measured by using ThermoFisher Quant-iT reagent and Qubit instrument and then diluted total RNA of each sample to a 50 ng/µL working concentration. RNA integrity was verified by electrophoretic profiling with Agilent Bioanalyzer 2100 (Agilent, CA) (Table S1).

### mRNA library preparation and RNA sequencing

To correct for the effect of the large amount of 5S rRNA on total RNA quantification in ovary samples, we measured the proportion of 5S rRNA peak area through Bioanalyzer electrophoresis using package *bioanalyzeR* [32] and then calculated the amount of total RNA required to meet the input (Table S1). 200 ng total RNA was used for RNA-seq library preparation using Illumina stranded mRNA Preparation Ligation kit. The 98 libraries were sequenced at the University of Helsinki Institute for Molecular Medicine Finland (FIMM) using an Illumina Novaseq 6000 sequencing platform producing 8,1 billion individual 100-bp paired-end reads.

### Quality control, alignment and assembly

RNA-seq reads were trimmed using fastp v0.23.2. [33] with default parameters and quality assessment of trimmed reads was performed using FastQC software. High-quality reads (Q>30) were mapped to the Atlantic salmon reference assembly Ssal_v3.1 (GCA_905237065.2) with Ensembl annotation release [34] (Salmo_salar-GCA_905237065.2-2021_07-genes.gtf) by employing STAR v2.7.11a [35] two-pass mode and the following steps: (i) 1-pass mapping: aligning the reads to the reference assembly, using mostly default parameters including: ( *--runMode alignReads –twopassMode None —-outFilterIntronMotifs RemoveNoncanonicalUnannotated --quantMode GeneCounts --alignIntronMin 20 --alignIntronMax 1000000 --alignSJDBoverhangMin 1 --alignMatesGapMax 1000000 --alignEndsProtrude 10 ConcordantPair --outFilterType BySJout --chimSegmentMin 10 --outSAMtype BAM SortedByCoordinate*); (ii) re-build STAR genome index: using Ensembl annotation and 1-pass splice junctions with parameters: (*--runMode genomeGenerate --sjdbFileChrStartEnd /path/to/sjdbFile.txt --limitSjdbInsertNsj 6000000 --sjdbOverhang 100 --genomeChrBinNbits 18* ); (iii) 2-pass mapping: mapping RNA-seq reads to the reference genome, parameter were (*--runMode alignReads –twopassMode None --outFilterIntronMotifs RemoveNoncanonicalUnannotated --quantMode GeneCounts --alignIntronMin 20 --alignIntronMax 1000000 --alignSJDBoverhangMin 1 --alignMatesGapMax 1000000 --alignEndsProtrude 10 ConcordantPair --outFilterType BySJout --chimSegmentMin 10 --outSAMtype BAM SortedByCoordinate --limitSjdbInsertNsj 6000000 --sjdbFileChrStartEnd /path/to/sjdbFile.txt*) (see Supplementary File 1 ).

The aligned read data were assembled into transcripts, including annotated and unannotated ones, using StringTie v2.2.1 with default parameters [36] with the reference genome Ssal_v3.1 and Ensembl annotation for a reference-guided assembly. After assembling the reads from all 98 samples separately, we applied StringTie *--merge* function to merge predicted transcripts from all libraries into a unified transcriptome.

The merged transcriptomes were passed to gffcompare v0.12.6 [37] for comparing with the reference annotation, to determine how many assembled transcripts were completely matched with annotated gene structures, in addition to identifying novel transcripts (class code “u”). The sensitivity and precision of the unified transcript are 99.5% and 36.1% respectively.

To benchmark the intergenic transcripts identified in our study, we reanalyzed two publicly available Atlantic salmon testis transcriptome datasets (PRJNA380580 and PRJNA550414) [25, 26] using our analysis pipeline. For this benchmarking, we indexed the Ssal_v3.1 genome using the unified annotation derived from our meta-transcriptome, which includes both Ensembl reference annotations and newly assembled transcripts. The testis datasets were then processed using STAR v2.7.11a in 1-pass alignment mode, followed by transcript assembly with StringTie v2.2.1. Finally, transcript structures were compared to both Ensembl reference annotation and the unified annotation using gffcompare v0.12.6 to assess the overlap and novelty of transcripts detected in the independent testis datasets.

### Long non-coding RNA identification

Traditional approaches to identifying lncRNAs through exclusion criteria are prone to false-positive predictions. Thus, we applied an approach that uses machine learning to predict lncRNAs, which improves the discrimination between lncRNAs and coding genes [38, 39]. We applied the lncRNA prediction tool FEELnc [39] to identify lncRNAs within the newly established merged annotation file of the pituitary-gonadal transcriptome in the 98 samples. First the FEELnc filter was used to filter and remove protein-coding, pseudogene, miRNA etc. and capture monoexonic transcripts with a minimal size 200bp. Second, using the main pipeline FEELnc codpot to compute the coding potential score for each of the candidate transcripts. Since the absence of species-specific lncRNAs set for Atlantic salmon, a machine-learning approach was used to simulate non-coding RNA sequences to train the model. We applied a “*shuffle*” strategy to take and shuffle the set of mRNAs while preserving 7-mer frequencies using Ushuffle. To calculate the coding potential score (CPS) cutoff separating coding (mRNAs) versus long non-coding RNAs (lncRNAs) (Figure S1), FEELnc codpot uses a R script that will make a 10-fold cross-validation on the input training files and finally, extracts the CPS that maximizes sensitivity and specificity. Finally, the FEElnc classifier applies a sliding window strategy (size: 10,000 to 100,000 bp) to classify lncRNAs based on their genomic localization with respect to nearest transcripts (protein-coding, non-coding or other loci). In addition, gffcompare [37] was applied to evaluate if the putative lncRNA transcripts detected from this merged transcriptome were intergenic by comparing to the Ssal_v3.1 annotation with class code “u” assignment considered as the indication of long intergenic non-coding RNA (lincRNA).

### Protein coding gene prediction from intergenic transcripts

We applied TransDecoder v5.7.1 [40] to identify candidate coding regions within the 12,383 nucleotide sequences of the class-U transcripts. First, open reading frames (ORFs) of at least 100 amino acids long were identified from the class-U transcript sequences. Second, to further enhance sensitivity for capturing ORFs with potential functional significance, we scanned all ORFs for homology to known proteins to identify common protein domains by searching the Pfam database (release 33.1) [41] using HMMER v3.3.2 [42]. Finally, ORFs with a Pfam hit were retained in the final output, ensuring that genome-based coding region predictions included sequences with strong evidence of function coding potential.

The newly predicted protein sequences were submitted to EggNOG-mapper v2 [43] using default parameters for function annotation. This tool provides predictions for functional categories, including functional descriptions and Gene Ontology (GO) terms, by mapping sequences to orthologous groups with known functions [43, 44]. Manual curation of selected candidate newly characterized protein-coding genes and lincRNA of interest was performed by using the IGV v2.19.1 [45] to visually inspect their genomic context and transcriptional evidence.

### Read count and differential gene expression analysis

We used featureCounts (subread-v2.0.6) [46] with the newly merged annotation file to calculate read count matrices for all 98 samples with all annotated loci. We used the count reads for performing DE analysis with the DESeq2 R package [47]. Before conducting differential expression (DE) analysis of the samples, we filtered the genes to exclude low-expressed loci by requiring that counts be greater than three in at least two samples. Then, the count data were normalized using the default *estimateSizeFactors* method in DESeq2 to account for differences in sequencing depth among samples. The normalized count data were further transformed with a *variance-stabilizing transformation* (VST) to facilitate Principal Component Analysis (PCA) for quality control and to assess the overall transcriptomic variation among the samples (Figure 3A).

DESeq2 was run with the design formular: *design = ∼ tissue* for investigating the tissue effects on gene expression. Each of the four tissues was compared with the others (in total, six pairwise comparisons) to identify significantly differentially expressed loci (protein coding and non-coding) using the Wald test, and p-value were adjusted using the Benjamini-Hochberg method to control the false discovery rate (FDR) [48]. We applied log2 (fold-change) ≥ 2 and adjusted P ≤ 0.01 to identify significantly upregulated loci. (see Supplementary File 2).

### Assessment of tissue-specific expression

We normalized data using transcripts per million (TPM) by ‘*Expression Calculator*’ function of TBtools v0.665 [49] for tissue-specific screening and the tissue-specificity index *τ* was evaluated as previously described [50]. The following criteria were applied to identify tissue-specific loci:

1. A locus with TPM < 1 in a given sample was considered as a not expressed locus and these TPM values were set to 0, we used the log_2_ transformed TPM;
2. A mean value from all replicates for each TPM is calculated of each locus;
3. Loci not expressed in any tissue were removed;
4. A tissue-specificity index *τ* for each gene was calculated by using the formula:

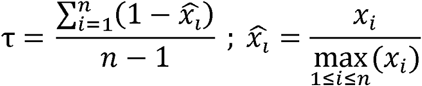

where *n* is the number of tissues and *x_i_* is the expression of the locus in tissue *i*

To call loci tissue specific, a threshold of 0.8 was set as previously recommended [51].

### Weighted Gene Co-expression Network Construction and Gene Module Detection

Weighted gene co-expression network analysis (WGCNA) was conducted using the R package WGCNA v1.72 . Gene-level counts were transformed using the ‘*vst*’ function from DESeq2, and only loci (protein-coding genes and lncRNAs) with expression variance ranked in the top 75 percentile of the data set were retained to reduce noise [52, 53].

To construct the co-expression network, we computed an adjacency matrix based on pairwise correlations between genes and applied a soft thresholding power of 16, which was selected based on the scale-free topology fit index (R^2^ > 0.8) (Figure S2). To capture biologically meaningful inverse relationships, such as those between lncRNAs and their targets [54], we applied an unsigned weighted network to identify protein-coding gene and lncRNA co-expression across tissues. A Topological Overlap Matrix (TOM) was then generated to capture network connectivity, and genes were clustered using hierarchical clustering (Figure S3). Gene modules were identified using the ‘*blockwiseModules*’ function in WGCNA, with the following parameters: ‘*TOMType = “unsigned”, minModuleSize = 30, maxBlockSize = 10000, deepSplit = 2.5, reassignThreshold = 0, and mergeCutHeight = 0.25*’.

For correlation analysis, tissue categories were binarized with the ‘*binarizeCategoricalVariable*’ function of WGCNA package with option ‘*includeLevelVsAll = TRUE*’, then we applied ‘*cor*’ function of WGCNA with ‘*use = “p”*’ to calculate the associations between module eigengene (ME) and tissue group by Pearson correlation. Those modules that were highly significantly correlated with at least one tissue (r>|0.7| and p < 0.01).

### Function Enrichment Analysis

For a comprehensive view, we generated the enrichment background gene set using known and newly characterized genes. We used the gProfiler [55] with ‘*g:Convert*’ function to fetch the gene annotations including GO terms for known genes combined with eggNOGmapper annotation for the newly characterized genes. To illustrate the functions of the gene cluster, we performed Gene Ontology (GO) enrichment analysis using R package clusterProfiler v4.12 [56], for the known and newly characterized protein-coding genes that were identified in each module. We selected the 10 most significantly enriched pathways to visualize in the maps, using a Benjamini & Hochberg false discovery rate (FDR) < 0.05 to identify significantly enriched GO terms.

## Results

### Large salmon RNA-seq dataset and bioinformatic workflow

We sequenced polyadenylated RNA from 98 pituitary and gonad tissue samples, producing a total of 8.1 billion paired-end 100 base pair raw reads with an average depth of 82 million reads per library, and a Q30 score above 93% (Table S2). After trimming of adapters and removing low-quality sequences, 8 billion clean reads were mapped to the Atlantic salmon reference genome Ssal_v3.1 with 72.6% uniquely mapping efficiency (Table 1). To better understand the molecular mechanisms underlying sexual maturation in Atlantic salmon, we developed a comprehensive bioinformatics pipeline to create an RNA-seq atlas for salmon gonad and pituitary tissues (Figure 1). We produced align-then-assemble transcriptome assemblies for each of the samples separately. The resulting 98 assembled transcriptome annotations were merged into one meta-assembly resulting in 502,232 transcripts (96,473 loci) with 5.2 transcripts per locus. This is significantly more than in the Ssal_v3.1 reference annotation that includes 181,901 mRNAs in 66,765 loci (∼2.7 transcripts per locus).

**Figure 1.**
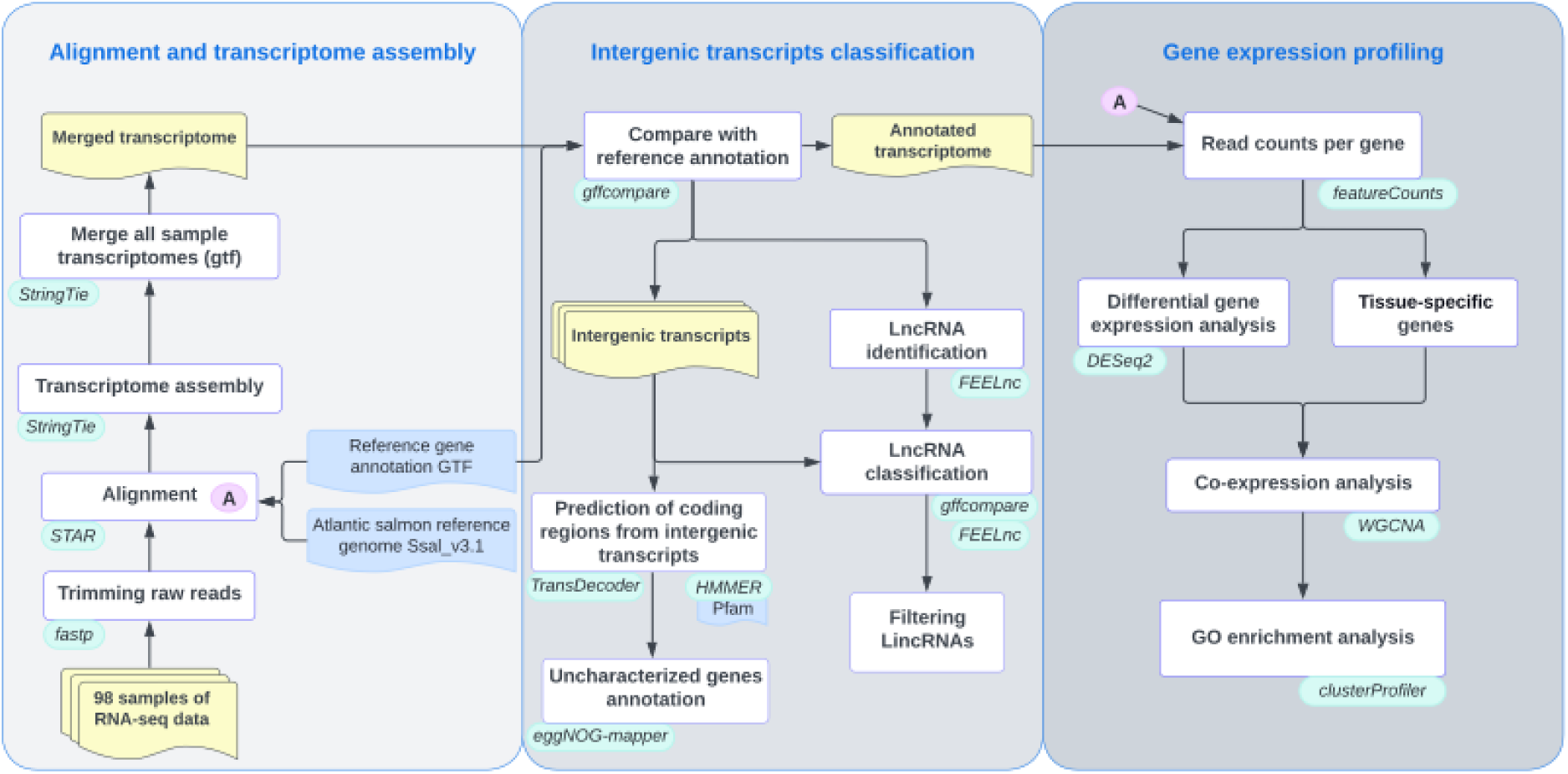
Overview of bioinformatic workflow showing the different steps performed to detect newly characterized protein-coding genes and long non-coding RNAs (lncRNAs) in the Atlantic salmon gonad and pituitary transcriptomes and to predict their potential functional role using gene co-expression and pathway enrichment analysis. Analysis workflow describing the data (yellow), software (green), reference data (blue), process (white).

**Table 1.** Average alignment results of transcriptome data for each tissue type. All values are in millions of reads.

|  | 1-pass mapping |  |  |  | 2-pass mapping |  |  |
| --- | --- | --- | --- | --- | --- | --- | --- |
|  | Total reads | Total map | Unique map | Multiple map | Total map | Unique map | Multiple map |
| Mature testis | 28.1 | 22.5 | 19.1 (67.2%) | 2.8 (10.4%) | 22 | 18.6 (65.3%) | 3.1 (11.5%) |
| Immature testis | 36.6 | 30.5 | 27.1 (73.4%) | 3.0 (8.2%) | 30 | 26.6 (72%) | 3.4 (9.2%) |
| Ovary | 46.2 | 38.5 | 32.3 (69.2%) | 4.6 (9.9%) | 37.6 | 31.3 (67.1%) | 5.0 (10.8%) |
| Pituitary | 29.7 | 26.1 | 23.6 (78.8%) | 2.1 (7.2%) | 26 | 22.7(76.1%) | 2.8 (9.7%) |

### Identification of protein-coding regions from intergenic RNA transcripts

To categorize transcripts, we compared the genomic coordinates of the assembled transcripts against the reference annotation using gffCompare [37]. Among the 502,232 transcripts, 36.1% (181,438) were classified as precisely matching known transcripts (class “=”), and 35.6% (178,974) of the transcripts comprised of multi-exon transcripts with at least one known exon junction (“j”). Interestingly, 6.3% (31,722) of the transcripts were classified as intergenic transcripts (“u”), representing transcribed regions that do not overlap with any annotated gene or pseudogene (Figure 2). Chromosomes 9,2 and 1 contained the highest number of intergenic transcripts (Figure S4). To assess the reliability and tissue consistency of the identified intergenic transcripts, we evaluated their representation across samples (Table S3). Overall, 83.7% (26,566) of the intergenic transcripts were supported by at least one sample, and 71.7% (22,760) were present in more than five samples, indicating high reproducibility (Figure S5). 1,752 intergenic transcripts were exclusively detected in a single tissue and were supported by all samples from that tissue, suggesting potential tissue specificity (Table S3, Figure S6).

**Figure 2.**
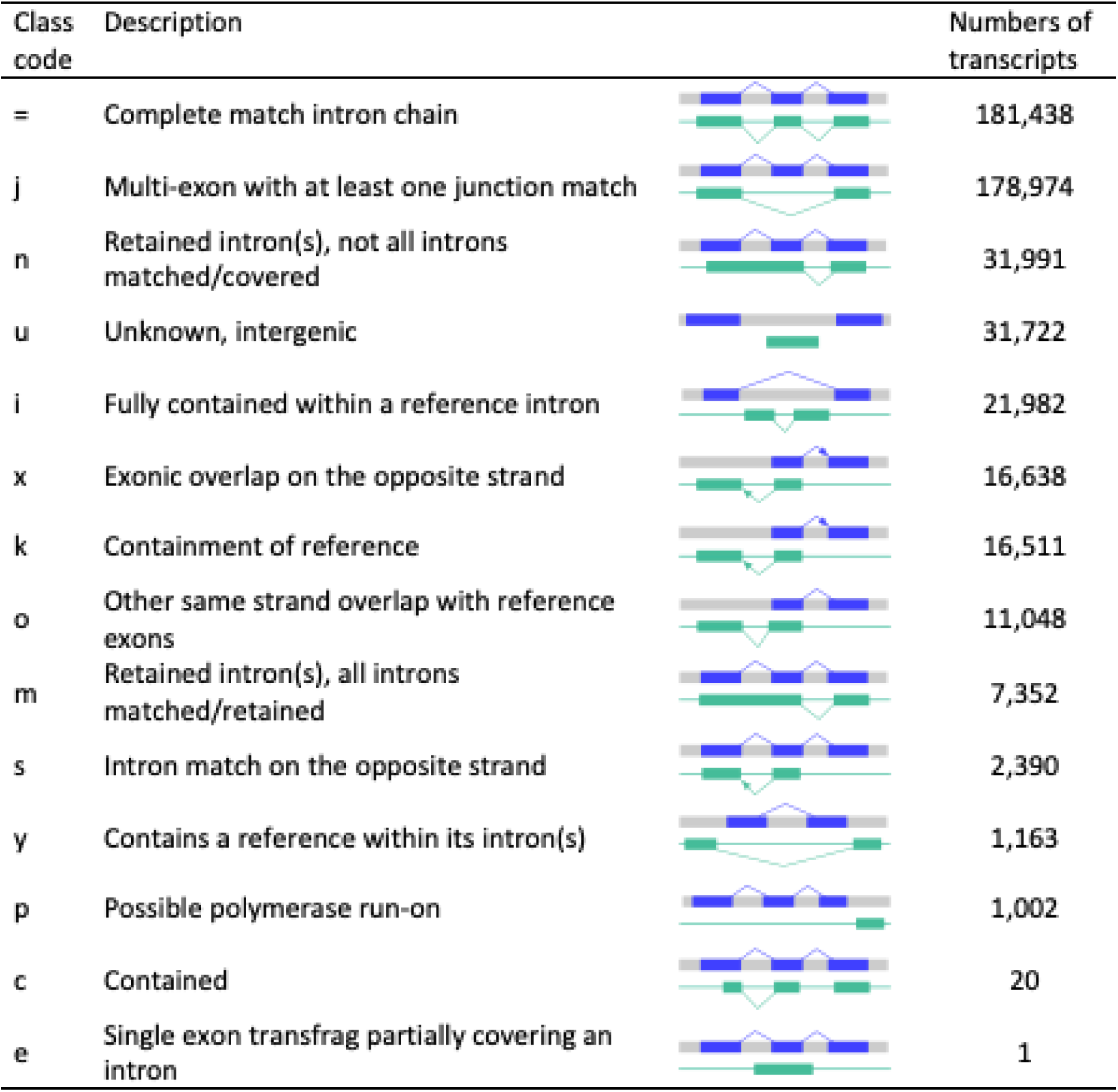
Number of transcripts in each transcript-classification code defined by gffCompare

Benchmarking to other existing salmon testis RNA-seq datasets PRJNA380580 and PRJNA550414 [25, 26] (Table S4-S6), our meta-assembly allowed us to identify 14,750 (on average 7.6% of all transcripts per samples) and 14,566 (6.8%) intergenic transcripts, with 79.6% and 70.7% detected in at least three samples, supporting their reproducibility and biological relevance (Figure S7, S8). Of these, 89.1% and 85.3% were complete or partial matches to intergenic transcripts in our meta-assembly intergenic transcripts, indicating that our assembly strategy provided a robust way of identifying previously uncharacterized transcripts. To note is that 37.0% and 40.1% of the matched intergenic transcripts were shorter than their counterparts in our data, suggesting more complete reconstruction. The remaining 8.8% and 12.4% unmatched intergenic transcripts may represent true biological differences between datasets.

To identify protein-coding regions from intergenic transcripts, we used methods that predict coding regions based on sequence homology to known protein-coding genes. Of the intergenic transcripts, 51.4% (16,305/31,722) encoded proteins longer than 100 amino acids. Functional annotation using EggNOG-mapper identified 6,100 intergenic transcripts associated with orthology terms.

### Identification and Characterization of LncRNA

We used a strand specific, directional library preparation for polyadenylated RNA with rRNA removal to enrich non-rRNA transcripts, making our data highly suitable for lncRNA discovery and analysis. First, we identified 29,544 lncRNA loci (73,825 lncRNA transcripts) by using FEELnc to annotate lncRNAs from RNA-seq assembled transcripts. This corresponds to 2.5 lncRNA transcripts per lncRNA locus, which is notably lower than the average number of transcript isoforms across all gene loci. We found an equal distribution of these lncRNA loci on both strands, with 49.8% (14,713) identified on the plus strand and 50.2 % (14,831) on the minus strand.

Moreover, lncRNAs can be categorized in several sub-classes based on their genomic position, and structure [57] i.e., genic lncRNAs, long intergenic noncoding RNAs (lincRNAs), anti-sense lncRNAs and sense lncRNAs. Understanding the relationships between newly annotated lncRNA transcripts and their closest annotated transcripts, especially in identifying potential lncRNA/mRNA pairs, is essential to for guiding functional annotations for lncRNAs [39, 58]. For this purpose, we applied FEELnc classifier module which employs a sliding window strategy around each lncRNA to capture all annotated transcripts located within the window. Out of 29,544 candidate lncRNA loci, 27,996 (71,086 lncRNA transcripts) had a nearby annotated partner locus (mRNAs).

Next, we identified 17,023 lincRNA transcripts from total 73,825 lncRNA transcripts using gffCompare. LincRNAs, defined as transcribing non-coding RNAs longer than 200 nucleotides that do not overlap annotated coding genes or pseudogenes, are known to play crucial roles in gene regulation, including gene expression control, scaffold formation and epigenetic control [59].

### Differentially expressed loci

To explore the transcriptome variation across tissues related to sexual maturation, we estimated gene expression levels from the assembled transcriptomes. Expression was estimated as read counts for all samples across 94,918 characterized loci, including (1) known protein-coding genes annotated in the reference genome, (2) newly characterized protein-coding genes within intergenic regions, (3) known lncRNAs annotated in the reference genome, and (4) lincRNAs newly identified within intergenic regions. We defined newly characterized protein-coding genes as those comprising solely of intergenic transcripts with coding capacity, while lincRNAs were defined as the lncRNAs consisting solely of intergenic transcripts without coding capacity, resulting in a total of 2,364 newly characterized protein-coding genes and 4,421 lincRNAs.

We summarized the overall expression profiles of the samples using principal component analysis (PCA), which revealed distinct tissue type clusters (Figure 3A). Two outlier samples from the mature testis were excluded from further analysis to ensure accuracy, one likely due to a mislabeling in maturation stage and the other located between the ovary and mature testis clusters, suggesting possible sample contamination.

**Figure 3.**
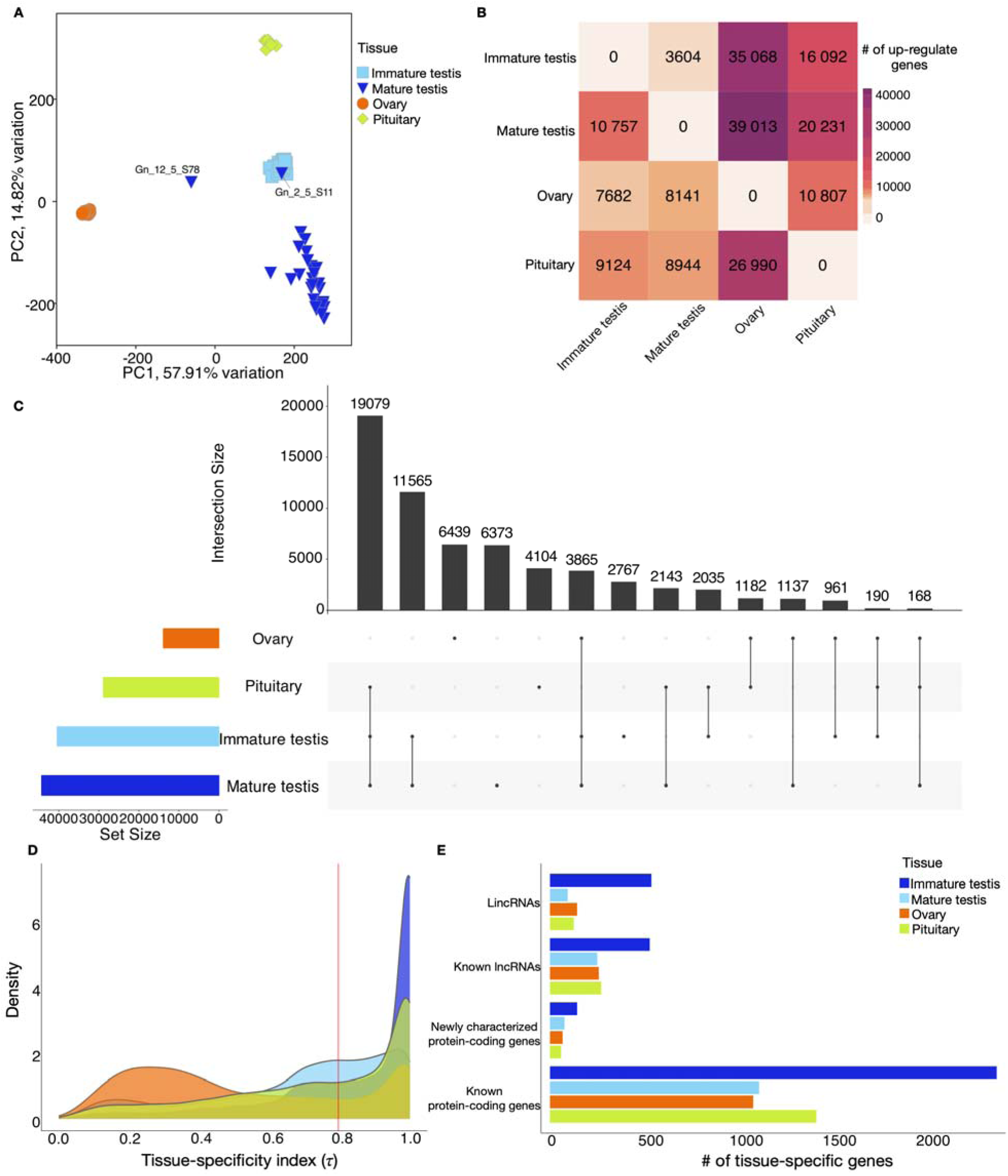
Overview of differential expression and tissue-specific loci across four Atlantic salmon tissues. (A), PCA performed with a normalized read count matrix of Atlantic salmon pituitary and gonad RNA-seq datasets with 98 samples (B), Heatmap of the numbers of upregulated genes in pairwise comparisons from the rows when compared with those from the columns. Gene upregulation was determined by using (log_2_(FC) ≥ 2 and padj ≤ 0.01) (C), UpSet plot showing the intersections among the upregulated genes across four tissue types. The histogram on the bottom left-hand side represents the total number of upregulated genes per tissue, the plot on the top right-hand side represents the number of upregulated genes that met each unique combination. The top barplot represents the sizes of the intersections between these different sets (D), A density plot of tissue-specificity index (*τ*) identified for each gene, based on average transcripts per million (TPM) in each tissue. Red line is the threshold (*τ* = 0.8) set for distinguishing tissue-specific genes (E), Barplot showing the number of tissue-specific genes identified across four tissues categorized into previously known/newly characterized protein-coding genes/known lncRNAs/ lincRNAs.

A pairwise comparison of global expression patterns across the four tissue types (Figure 3B) identified 62,008 loci (65.3%) that were differentially expressed loci (DEL; |log_2_(FoldChange)| ≥ 2, adjusted *P* ≤ 0.01) in at least one tissue compared to other tissues. The number of DEL varied widely between tissue comparisons, ranging from 14,361 in mature vs. immature testis comparison (10,757 upregulated, 3604 downregulated) to 47,154 in mature testis vs. ovary comparison (39,013 upregulated, 8,141 downregulated). Moreover, comparisons with the ovary yielded large numbers of upregulated loci, with 35,068, and 26,990 upregulated in immature testis and pituitary respectively, suggesting generally lower expression levels in immature ovary.

In general, mature testis comparisons yielded the highest number of upregulated loci, followed by immature testis, pituitary, and ovary (Figure 3C). The largest intersection of upregulated loci (19,079) was shared among pituitary, immature testis, and mature testis when compared to ovary. Both ovary and mature testis had similar numbers of unique upregulated loci, with 6,457 and 6,345 respectively. Overall, we identified 19,683 loci (ovary: 6,439; mature testis: 6,373; pituitary: 4,104; immature testis: 2,767) as upregulated in a single tissue compared with all the other tissues, indicating tissue specificity.

### Tissue specificity in gonad and pituitary

Quantification of tissue specificity is a powerful statistical analysis approach to investigate the correlation between tissue-specific expression and development, helping to define marker genes for each tissue type and to reveal tissue-specific molecular regulation [60, 61]. To do this, we normalized data using TPM and employed the tissue-specificity index *τ* (Tau) to determine whether a gene is tissue specific or broadly expressed [50]. Of the 94,918 loci analyzed, 82.2% (78,057) were expressed (TPM>1) in at least one sample, with the highest numbers in mature testis (70,458), immature testis (66,176), pituitary (51,097), and ovary (37,768) (Figure S9). Conversely, 16,662 loci were not expressed in any tissue, with 4,776 loci not expressed in any sample.

A total of 37,355 loci (39.4%) were assigned *τ* > 0.8, indicating strong preferential expression in specific tissues. However, the distribution of these genes varied greatly among different tissue types (Figure 3D). Mature testis exhibited *τ* distributions with largest peaks near 1, suggesting strongest tissue-specific expression, followed by immature testis and pituitary. From differential expression analysis, 19,683 loci identified as upregulated in a single tissue, and 52.6% (10,349) of these had *τ* values greater than 0.8. Based on their strong preferential expression in a particular tissue, these 10,349 loci were classified as tissue-specific loci in this study. Mature testis had the highest number of tissue-specific loci (4,449), followed by pituitary (2,192), immature testis (1,899) and ovary (1,809). Only 27.6% (1,809/6439) of loci upregulated in only ovary were determined as tissue-specific loci. In contrast, the percentages for mature and immature testis were significantly higher, at 69.8% (4,449 out of 6,373) and 68.6% (1,899 out of 2,767), respectively, indicating that while a large number of loci were upregulated in the ovary compared to other tissues, expression of most of these loci was not specific to ovary.

Categorizing these tissue-specific loci into known and newly characterized protein-coding genes or known/newly characterized lncRNAs and lincRNAs showed that mature testis contained the highest number in all four categories (Figure 3E). While the overall expression level of newly characterized protein-coding genes was higher than that of lincRNAs (Figure S10), lincRNAs were more frequently tissue-specific. Mature testis contributed more than half of all tissue-specific intergenic loci (both newly characterized protein-coding genes and lincRNAs), followed by the pituitary. The ratio of lincRNAs/lncRNAs (914/1,327) was markedly higher for newly characterized gene to previously known genes (352/6,047) (Figure 3E), indicating that our analyses had identified previously uncharacterized transcriptome variation especially when considering lncRNAs. For example, there were more lincRNA (542) than known lncRNAs (526) in the mature testis transcriptomes. These ratios were lower for immature testis (95/256), ovary (148/263) and pituitary (128/277), but still considerably higher than the ratio of newly characterized to previously known protein-coding genes in these tissues (mature testis: 146/2,339; immature testis: 80/1,180; ovary: 68/1,120; pituitary: 60/1,440).

### Co-expression network of lncRNAs with protein-coding genes

Weighted gene co-expression network analysis (WGCNA) is a common and powerful approach for revealing meaningful relationships between gene modules and biological processes, and between genes and traits [62–64]; we used WGCNA to identify protein-coding gene and lncRNA co-expression network modules. By clustering correlated genes together, we identified nine distinct gene network modules with the number of loci per module ranging from 35 in the pink module to 6,433 in the turquoise module (Figure 4B). Protein-coding genes dominated each module, especially in large modules such as turquoise (5,239), brown (4,580), blue (4,543), yellow (3,949) (Figure 4B). Newly characterized protein-coding genes and lincRNAs were more abundant in the turquoise (240) and blue modules (183), while lncRNAs were found across all modules but were particularly high in turquoise (495), blue (451), and brown (362) (Figure 4D).

**Figure 4.**
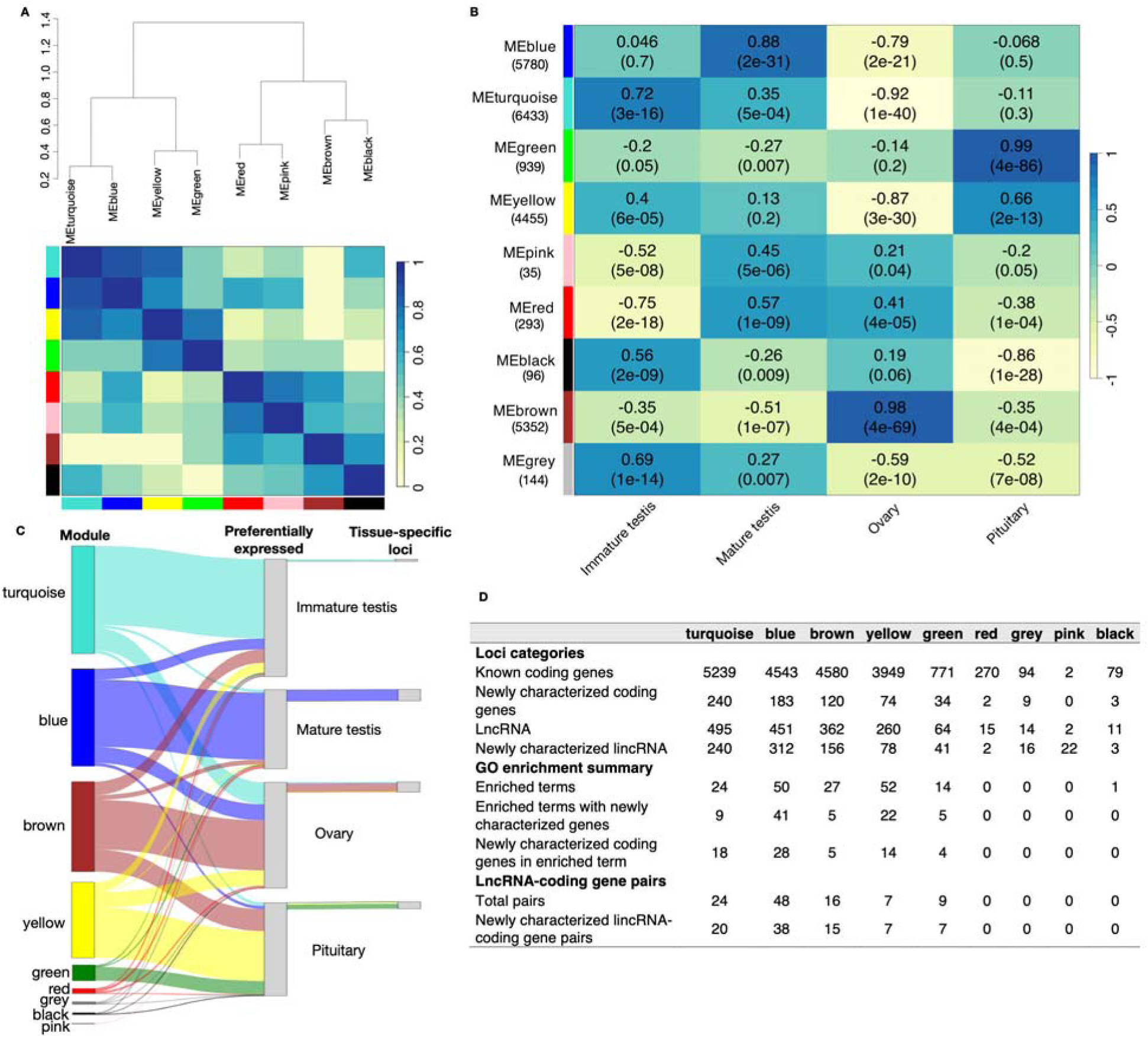
Identification of gene modules associated with Atlantic salmon reproductive tissues. (A), Eigengene dendrogram and heatmap showing the correlation among nine modules. The upper panel presents a hierarchical clustering dendrogram of module eigengenes (labeled by module colors). The lower heatmap displays the eigengene network, where each row and column in the heatmap corresponds to a module eigengene (labeled by module colors), in the heatmap, blue represents low adjacency (negative correlation) while red represents high adjacency (positive correlation). (B), Heatmap showing the relationships of module eigengene and tissues. Each column corresponds to a tissue, and each row corresponds to a module eigengene, module names are indicated on the left with the gene number in parentheses. Pearson’s correlation coefficient is used as the correlation descriptor, with blue and red indicating negative and positive correlations, respectively, and p-values are shown in brackets. (C), Sankey diagram illustrating the numbers of loci in each module and their preferential expressed tissues and tissue specificity. The first column represents the number of loci of each module (labeled by module colors). The second column represents the number of loci preferentially expressed in each tissue, determined based on mean TPM values. The third column represents the number of tissue-specific loci. The connection between terms is illustrated according to the distribution of the term and module color. (D), Table showing an overview of each module including the loci type distributions, GO enrichment terms, and interaction between lncRNAs and coding genes.

We first summarized the relationships among the modules and found that module eigengenes exhibited highly significant correlations, e.g. the turquoise and blue and yellow and green modules are highly correlated (Figure 4A). Further, we investigated the associations of expression profiles between each module and each tissue using Pearson correlation analysis, where each tissue was binarized and tested against all other tissues. This analysis aimed to determine whether each module’s expression profile varied significantly across the four tissues. We found four modules that were strongly positive correlated with single tissues at r > 0.7 and p < 0.01 (Figure 4B). The green module was strongly correlated with pituitary (r=0.99), the brown module was strongly correlated with ovary (r=0.98), and the blue and turquoise modules were correlated with mature and immature testis respectively (r=0.88; r=0.72). By comparing the loci in each module to the tissue specific loci for each tissue (defined as comparison group), we found that the tissue specific loci and the loci in each module were highly consistent for each comparison group (Figure 4C). Most of the genes in the blue module were preferentially expressed in mature testis samples with 69% (3390/5780) loci showing high expression in mature testis. Among these, 679 were identified as mature testis specific loci. The turquoise module showed high expression in immature testis and some mature testis samples; 74% (4744/6433) loci were preferentially expressed in immature testis and 119 were immature testis specific loci. The green and yellow modules exhibited elevated expression in pituitary samples; 88% (823/939) loci clustered in green module were preferentially expressed in pituitary and 298 were pituitary specific loci. Additionally, 66% (2933/4455) loci clustered in yellow module were preferentially expressed in pituitary and 118 were pituitary specific loci. The brown module was prominently expressed in the ovary; 55% (2956/5352) of the loci were preferentially expressed in ovary and 499 were ovary specific loci.

### Gene ontology term enrichment analysis

We performed gene ontology (GO) analysis to explore the functions of known/newly characterized protein-coding genes in each module identified in WGCNA (Table S7, Figure 5, S11). This approach enabled functional predictions for newly characterized protein-coding genes based on their associations with well-annotated genes. The terms cell adhesion, immune response, extracellular matrix organization, and Wnt signaling were enriched in turquoise module which is strongly correlated with immature testis expression (Figure 5A). The blue module, which was highly expressed in mature testis, was enriched in processes such as lipid and heme binding, oxidoreductase activity, cytoskeleton organization, and cilium movement involved in cell motility (Figure 5B). The brown module, linked to ovary expression, was enriched in rRNA processing, ribosome biogenesis, DNA replication and repair, and transcriptional regulation (Figure 5C). The green module, associated with pituitary expression, was enriched in hormone activity, synaptic transmission, and ion channel function (Figure 5D).

**Figure 5.**
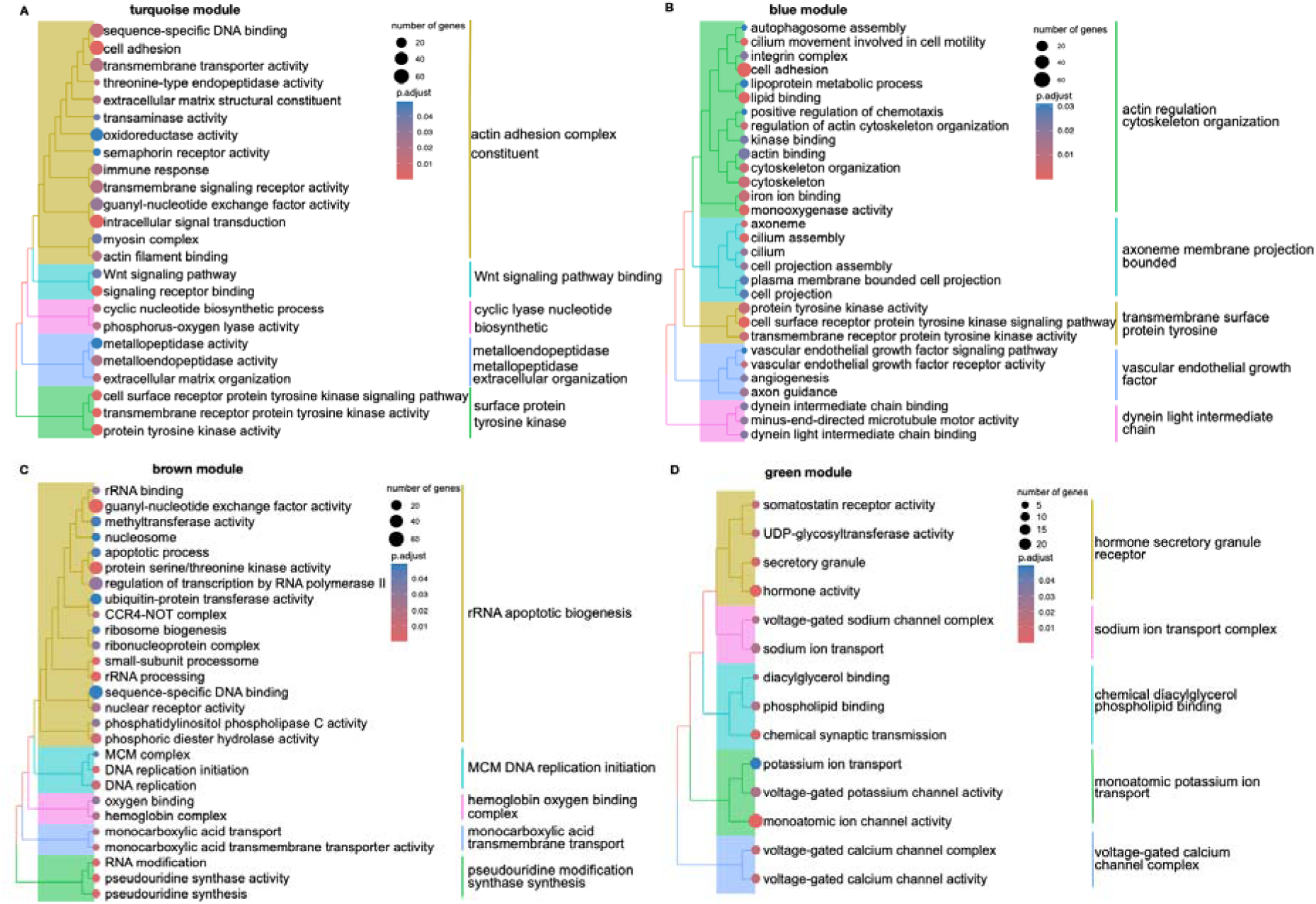
GO analysis of Atlantic salmon reproductive tissue-associated modules. Tree plots displaying significantly enriched GO terms for genes within WGCNA modules. The terminal branch of the tree represents a GO term, and the size of the dot shows the number of genes assigned from the module gene set to that term. The color represents the adjusted p value of the enrichment. The functionality that each cluster represents is denoted on the right side. (A) Immature testis module (turquoise): The top 30 significant enriched GO terms were clustered into five categories. (B) Mature testis module (blue): The top 30 significant enriched GO terms were clustered into five categories. (C) Ovary module (brown): The top 30 significant enriched GO terms were clustered into five categories. (D) Pituitary module (green): The top 14 significant enriched GO terms were clustered into five categories.

We also found 69 newly characterized protein-coding genes across four tissues within the enriched 82 GO terms (Figure 4D, Table S8), such as sequence-specific DNA binding, cell adhesion, microtubule, metallopeptidase activity, and secretory granule. Seven of these candidate newly characterized genes were identified as tissue-specific genes, five were mature testis-specific genes (such as *phc1, pop1,* and *hmgb1*), and two were pituitary-specific genes (*gh1* and *nbeal2*). Other candidate newly characterized genes including, *pou3f2, foxq1 mc1r* and *gjb3* in turquoise module *clip1, hmgb1* and *rab11fip5* in blue module, *ier5* and *anapc5* in brown module, and in green included *ugt8* and *amer2* (see more in Supplementary File3). Interestingly, most of these newly characterized genes appeared to have additional annotated copies located in different genomic regions. For example, *gh1* (MSTRG.66383) was identified on chromosome 6 in our RNA-seq data, while an annotated copy of *gh1* (ENSSSAG00000055013) already exists on chromosome 3. Both copies shared a similar expression pattern, being highly expressed in the pituitary. This pattern was also observed for other genes, such as *foxq1* paralogs were identified from the module, preferentially expressed in immature testis, *ier5* and its annotated paralog were found in the brown (ovary associated) module. This suggests that these newly characterized genes represent newly identified copies that exhibit similar expression profiles to their annotated counterparts.

### Functional prediction of lncRNAs (lincRNAs and NATs) based on co-expression with neighboring coding genes

LncRNAs can regulate the transcription of nearby genes (cis-regulation), particularly those located on the same or opposite DNA strand, or in intergenic regions [65, 66]. To predict lncRNA functions, we first used FEElnc to annotate lncRNAs by identifying their closest annotated genes or lncRNAs, generating candidate lncRNA-mRNA pairs. Subsequently, WGCNA grouped genes and lncRNAs into co-expression network module across four tissue types based on similar expression patterns. We prioritized lncRNA-gene pairs that were situated within 100k basepairs or each other and appeared in the same module, indicating not only genomic proximity but also shared expression patterns.

In total, we identified 104 candidate lncRNA-protein coding gene pairs across five major modules (turquoise, blue, green, brown and yellow) (Figure 4D, Table S9), with no pairs identified in the smaller modules. Of these, 46.2% (48 pairs) were from the blue module mostly highly expressed in mature testis (37 pairs), meanwhile 24 pairs were from the turquoise module, with 18 pairs highly expressed in immature testis, consistent with tissue-specific co-expression patterns observed in our earlier analysis. The remaining 32 pairs were distributed across the green, yellow, and brown modules. 83.7% (87/104) of these pairs involved newly characterized lincRNAs, while only 17 pairs included known lncRNAs, underscoring the importance of newly discovered transcripts in regulation.

The lncRNA annotation can provide hints for predicting lncRNA functions. For example, there were 11 lincRNAs annotated as ‘intergenic antisense upstream’ which correspond to divergent lincRNAs (i.e. transcribed in head to head orientation with the gene partner) among which five are less than 5kb from their gene partner transcription start sites (i.e. *tnfrsf11b, fgl1*) (Table 2). This class directly pinpoints to lincRNAs potentially sharing a bi-directional promoter with their gene partners [39, 67, 68]. On the other hand, 6 lincRNAs located less than 5kb from their gene partner, belong to the ‘sense intergenic upstream’ class and may correspond to dubious lncRNAs that are 5’UTR extensions of the neighboring protein-coding genes. Interestingly, based on tissue-specificity, we found that there were 13 partner protein-coding genes previously identified as tissue specific loci, 7 of which were uniquely expressed in mature testis, including *tafa3, lrfn1* and *fam46a*.

**Table 2.**
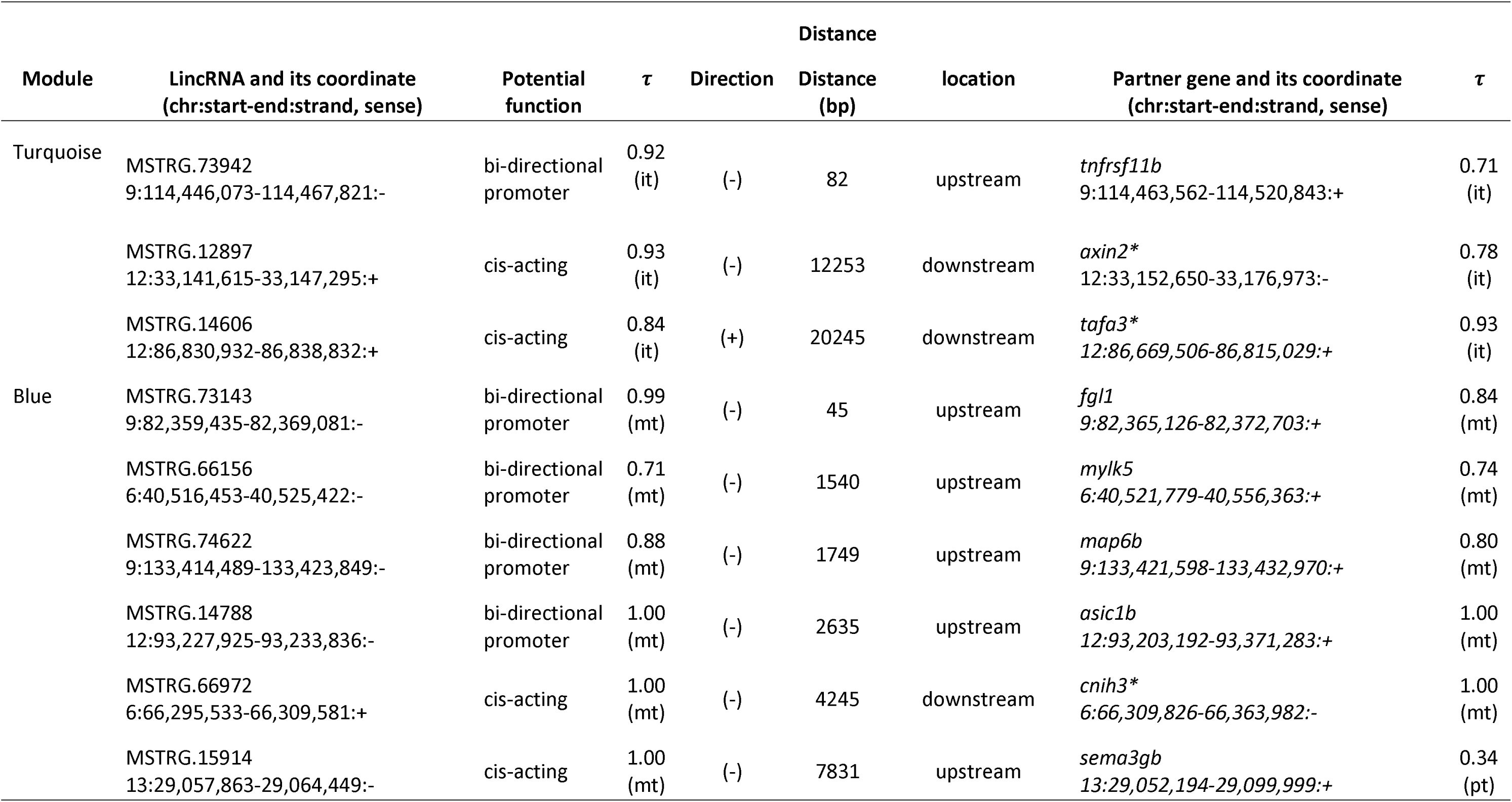
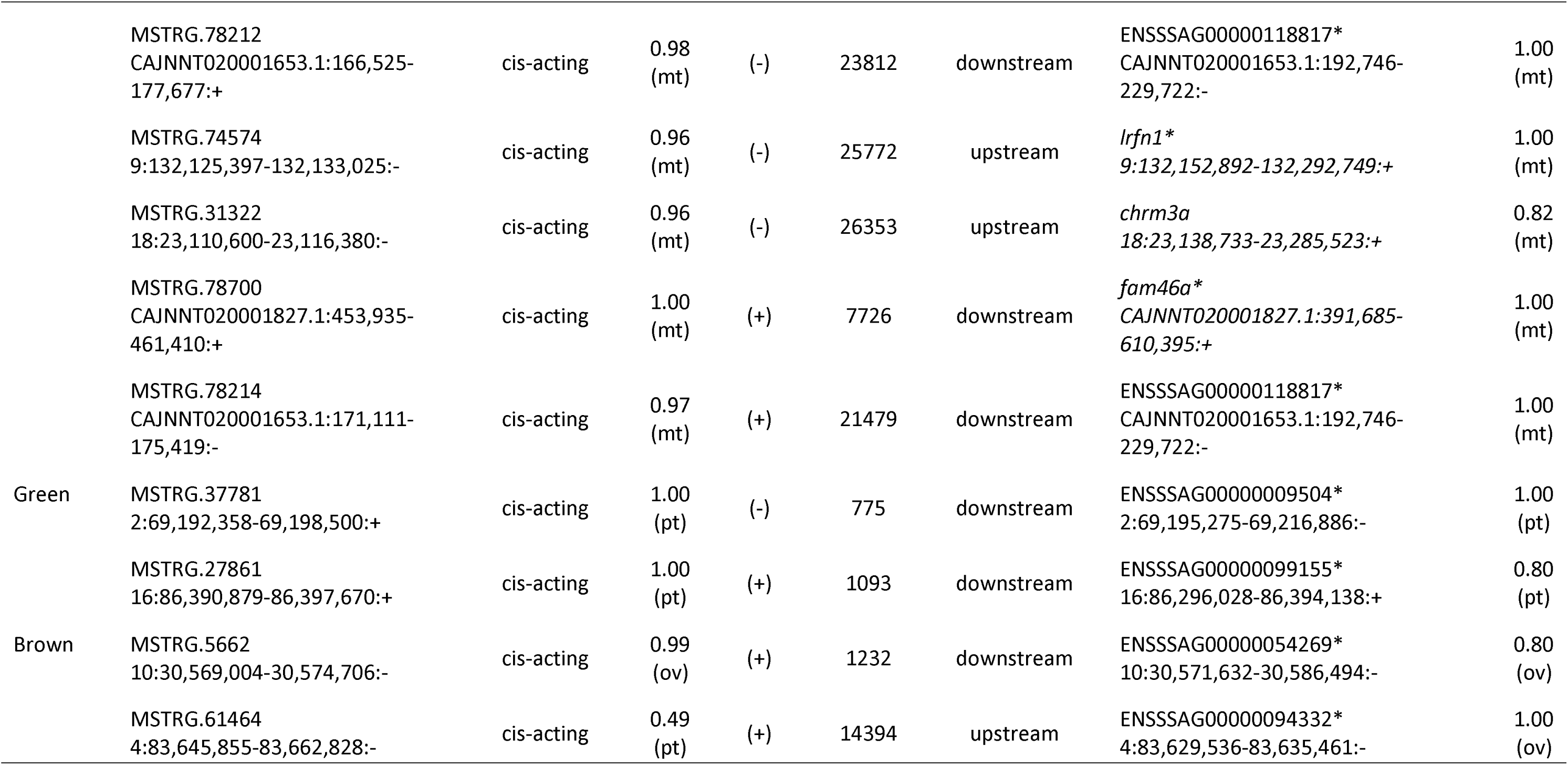
Potential functional lincRNAs. The (-) and (+) symbols indicate antisense and sense of the direction of lincRNA to its partner gene, *τ* value of each loci showing its tissue specificity * indicates loci upregulated in a single tissue, it: immature testis, mt: mature testis, ov: ovary, pt: pituitary

## Discussion

Understanding the complex mechanisms that control biological processes such as sexual maturation benefits from detailed exploration of the transcriptome. Despite advances in high-throughput sequencing technologies, current reference transcriptomes often capture only a fraction of the extensive RNA variation revealed by these methods, particularly for non-reference transcripts that may play critical roles in phenotypic traits [39, 67, 68]. Transcriptome analyses frequently rely on reductive identification methods, which can underestimate transcriptome complexity and exclude genes that are not previously characterized in specific tissues [23]. Our study aimed to address these gaps and succeeded in uncovering significant transcriptomic complexity in gonad and pituitary tissues of Atlantic salmon by identifying thousands of previously unreported genes and lncRNAs, offering novel insights into their potential functions.

### Transcriptomic complexity in gonads

Using a custom bioinformatic pipeline, we systematically identified and annotated previously uncharacterized (intergenic) protein-coding and lncRNA transcripts in salmon gonad and pituitary transcriptomes. While intergenic transcription has been commonly considered a consequence of pervasive transcription with no apparent function, accumulating evidence highlights its potential regulatory roles, including its influence on neighboring gene activity [69–71]. Some studies have reported that most de novo genes originate from intergenic regions [72] and exhibited testis-biased expression [72, 73]. We found that more than half of the intergenic transcripts had protein-coding capability (PCC), indicating that intergenic transcripts have potential importance of either extending transcriptional events for canonical protein-coding genes or to be as-yet unannotated genes. Alternative transcription start sites (ATSS) significantly contribute to isoform diversity, particularly in early developmental stages [74, 75], which could account the origin for some of the intergenic transcripts in this study. It is to note that some putative lincRNA transcripts were assigned to known genes after gene-level quantification, indicating some of them could be parts of mRNAs due to ATSS.

We found that the mature testis exhibited the most significant tissue-specific characteristics among the four tissues analyzed. This was evident in two key aspects: 1) differences in gene expression between mature testis and other tissues were most significant and 2) mature testis expressed the largest numbers of tissue-specific loci including known and newly characterized protein-coding gene copies, lncRNAs, and lincRNAs originating from intergenic regions. Overall, our study revealed widespread transcription and high tissue-specificity in mature testis, which has been previously reported in the literature for mammals and birds [76–78]. At the molecular and genomic levels, the testis is well established as the most rapidly evolving organ and particularly receptive to genomic innovations such as the emergence of new gene copies and structures [24, 79]. In contrast, immature ovary gene expression exhibited the lowest tissue-specificity, indicating the marked differences in gene expression levels between male and female gonads. This specificity is biologically relevant but previously uncharacterized in Atlantic salmon, providing new insights into regulation of tissue-specific transcription and the emergence of new gene copies in the salmon reproductive axis.

Several hypotheses have been proposed to explain widespread gene transcription in the testis. One explanation is that more genes are required for the functioning of testis [80], while another suggests the leaky transcription caused by massive chromatin remodeling during spermatogenesis [77, 81]. While the transcriptional scanning hypothesis has been proposed to explain the origin of widespread transcription in mammalian testis [82], some researchers argue that the transcriptional scanning cannot account for intergenic transcription, attributing it instead to transcription-associated mutagenesis [83]. Testing these hypotheses in future studies, particularly with respect to intergenic regions, could provide critical insights into the molecular mechanisms underlying sexual maturation in Atlantic salmon.

The annotation of the current Atlantic salmon reference genome included one sample each of pituitary, testis, and ovary. However, as shown in other animals [77, 81], our results suggest that the mature testis appears to be an excellent tissue for detection of protein-coding transcripts from either unknown genes or genes misannotated as non-coding. This highlights the limitation of relying solely on existing reference annotations for gonadal transcriptome studies. To address this, we recommend an align-then-assemble strategy [28], which allows for the identification of novel transcripts that may not be included in the reference annotation, improving the understanding of gene expression dynamics in reproductive tissues.

### Functional insights from tissue-specific gene network modules and newly characterized gene copies

We identified co-expression modules that were correlated with specific tissues. Gene Ontology (GO) enrichment analysis further demonstrated that the pathways associated with each tissue-specific module reflected the functional roles of these tissues. For example, wnt signaling was enriched in the module correlated with immature testis (turquoise). The wnt pathway is known for its roles in determining spermatogenic cell fate and maintaining spermatogonia [84–86]. Wnt signaling has been further shown to be altered during early puberty in fish like Atlantic salmon [27, 87]. Immune response processes in this module further reflected the establishment of immune privilege which is essential for protecting germ cells from the individual’s immune response [88].

We further identified 69 previously unknown protein coding gene copies with potential function in salmon maturation as they were associated with tissue-specific expression modules and over-represented GO terms. These genes are likely duplicated copies of previously annotated genes in the Atlantic salmon genome. This is consistent with the whole-genome duplication (WGD) in salmonid species, which led to a portion of the genome remaining in effective polyploidy [89, 90]. Interestingly, several of these newly annotated genes exhibited expression patterns similar to their annotated paralogs, suggesting that these duplicated genes may contribute to regulating salmon maturation. Our study highlights the importance of considering gene duplication when analyzing transcriptomic data in salmonids. Understanding the roles of these duplicated genes will be essential for future RNA-seq studies on Atlantic salmon maturation, as they may reveal novel regulatory mechanisms or compensatory pathways [91].

### Functional insights on the role of lncRNAs in sexual maturation

There is growing evidence that lncRNAs play important roles in many biological processes [92, 93] including reproduction [94]. Intergenic lncRNAs (lincRNAs) can act on nearby upstream or downstream genes via cis-regulation [95]. This regulatory relationship forms the basis for predicting the function of lncRNAs by examining their molecular interactions with neighboring genes and pathways.

In our analysis, all modules identified through WGCNA included both lncRNAs and protein-coding genes, suggesting coordinated regulatory roles. Many of the identified lncRNA-gene pairs showed physical proximity on chromosomes and shared co-expression patterns, reinforcing the possibility of cis-regulation. Interestingly, most candidate lncRNA-gene pairs involved lincRNAs rather than previously known lncRNAs, underscoring the need for their discovery to uncover new regulatory mechanisms. For example, *tnfrsf11b*, which was preferentially expressed in immature testis, paired with lincRNA MSTRG.73942 located in the opposite orientation 82bp upstream. The *tnfrsf11b* gene encodes for osteoprotegerin (OPG), which enhances sperm counts and male fertility by protecting testis from excessive inflammation or apoptosis [96]. Moreover, *fgl1*, which was specifically expressed in mature testis, paired with lincRNA MSTRG.73143 located 45bp upstream. *Fgl1* has been proposed to shield viable sperm from degenerating spermatozoa within the tubule [97], underscoring its role in sperm quality control. We also identified a lincRNA MSTRG.78700 at the downstream of *Fam46a* which has been shown to be down-regulated in rat Sertoli cells lacking endogenous androgen receptor [98]. *Fam46a* was furthermore identified as a mature testis-specific expressed gene in our analysis. Our characterization of these candidate lincRNAs supports the view that lincRNAs play important roles in the regulation of maturation processes. Additionally, the discovery of novel lincRNAs highlights the potential to uncover previously uncharacterized regulatory mechanisms between lncRNAs and protein-coding genes, although further experimental validation is needed to confirm these interactions.

### Limitations of reference-based transcriptomes

Most transcriptome studies of Atlantic salmon testis to date [13, 25–27] have successfully employed reference annotations to characterize gene expression. While this has provided valuable insights, the use of a reference model may not fully capture the complexity of the testis transcriptome, particularly in relation to novel or less well-annotated transcripts. When benchmarking to two existing salmon testis transcriptome datasets, we found over 14,000 intergenic transcripts that matched the newly characterized transcripts in this study, indicating that our conclusions are substantiated by comparative data and that our study fills a significant gap between the reference annotation and real transcriptome complexity. While experimental validation and cross-species analyses (e.g., orthology, synteny, or proteomics) would further enhance confidence in the functionality of the newly identified transcripts, our current annotations are informed by stringent computational criteria and a conservative, multi-layered filtering approach designed to minimize potential biological false positives. One important limitation of our study is the use of polyadenylated RNA enrichment for library preparation, which inherently biases the transcriptome toward polyadenylated transcripts and restricts the detection of non-polyadenylated lncRNAs [99, 100]. To achieve a more complete understanding of transcriptomic regulation, particularly in the testis, future studies could benefit from using ribosomal RNA depletion or total RNA-seq strategies that enable the detection of both polyadenylated and non-polyadenylated transcripts.

Our study further demonstrates that the choice of reference annotation can significantly impact RNA-seq read mapping and gene quantification [101]. Of the two major annotation databases, Ensembl offers broader gene coverage than RefSeq, which follows more stringent inclusion criteria. For example, 22,333 Atlantic salmon transcripts annotated in Ensembl were found in intergenic regions when compared against RefSeq, indicating that these would have been missed using RefSeq alone. In this study, we used the Ensembl annotation to maximize transcript discovery, particularly for intergenic transcripts potentially representing novel genes or isoforms. However, using a single annotation source may introduce biases and we generally recommend cross-referencing newly identified candidate genes with RefSeq annotations. Single-sample transcriptome reconstruction can further limit the downstream analysis across many samples [102]. Thus, our analysis generated a unified meta-assembly and uncovered thousands of intergenic transcripts. While such meta-assembly can introduce redundancy, we addressed this by assessing sample-level support for each intergenic transcript. Most intergenic transcripts were detected in multiple samples, indicating robust reproducibility. Furthermore, we performed gene-level quantification in downstream analysis which is a strategy helping to mitigate redundancy from the multi-sample transcriptome assemblies.

## Conclusion

This study provides a systematic and comprehensive transcriptome analysis of Atlantic salmon gonad and pituitary transcriptomes, significantly improving our understanding of the salmon reproductive transcriptome by identifying newly characterized putative protein-coding genes and lincRNAs from intergenic regions. By integrating differential expression analysis, tissue-specificity quantification, and gene network analysis, we have identified key regulatory pathways and potential functional lncRNAs that warrant further investigation. The findings expand the repertoire of annotated transcripts and reveal important tissue-specific expression patterns across four tissue types, with the mature testis showing particularly high tissue specificity especially in newly annotated genes and lncRNAs. These newly annotated genes and lncRNAs contribute valuable additions for Atlantic salmon genome annotation as well as insights into molecular mechanisms underlying sexual maturation and reproductive development, offering potential targets for further functional studies.

## Declarations

### Ethics approval and consent to participate

The rearing and sampling of Atlantic salmon were performed according to European animal welfare regulations and approved by the Finnish Project Authorization Board (permits ESAVI/2778/2018 and ESAVI/4511/2020).

### Consent for publication

Not applicable.

### Availability of data and material

Sequence data are publicly available in EMBL-EBI European Nucleotide Archive (ENA; https://www.ebi.ac.uk/ena/) under project number PRJEB86880.

Output files for transcriptome annotation and gene expression are publicly available in Zenodo repository (https://zenodo.org/records/15639686).

### Competing interests

The authors declare no competing interests.

### Funding

This research was funded by Academy of Finland (grant numbers 314254, 314255, 327255 and 342851), the University of Helsinki, and the European Research Council under the European Articles Union’s Horizon 2020 and Horizon Europe research and innovation programs (grant no. 742312 and 101054307), by the European Union under the Marie Sklodowska-Curie grant agreement no 10110362 (to J.-P.V.), and the China Scholarship Council (grant no. 202208530004 to X.-D.H.). Views and opinions expressed are however those of the author(s) only and do not necessarily reflect those of the European Union or the European Research Executive Agency. Neither the European Union nor the granting authority can be held responsible for them.

### Authors’ contributions

Conceptualization C.R.P., J.-P.V., X.-D.H., Formal analysis X.-D.H., Methodology X.-D.H., J.-P.V., M.F., C.R.P., Investigation X-D.H., J.-P.V., C.R.P., Resources C.R.P., J.-P.V., E.P.A., Supervision C.R.P., J.-P.V., Funding acquisition: C.R.P., J.-P.V., X.-D.H., Visualization: X.-D.H., Writing – original draft X-D.H., Writing – review & editing X.-D.H., J.-P.V., C.R.P., M.F., E.P.A.

## Acknowledgements

We thank Jaakko Erkinaro and the Natural Resources Institute, Finland (Luke) for access to broodstock material. We thank all the people for taking care of the fish in Viikki and Lammi. We thank FIMM Genomics unit for providing sequencing and CSC – IT Center for Science, Finland, for computational resources. We thank Nikolai Piavchenko for fish sampling. We thank Annukka Ruokolainen, Iikki Donner and Mikaela Hukkanen for RNA extraction and Katja Maamela, Nora Bergman and Gersende Maugars for useful discussions.

## Notes

### Competing Interest Statement

The authors have declared no competing interest.

