## Supplemental figure and table for "A Comprehensive Analysis of Atlantic Salmon Gonad and Pituitary Transcriptomes Identifies Novel Players in Sexual Maturation": ms_salmon_gnpt_rnaseq_Supplementary_figures.docx

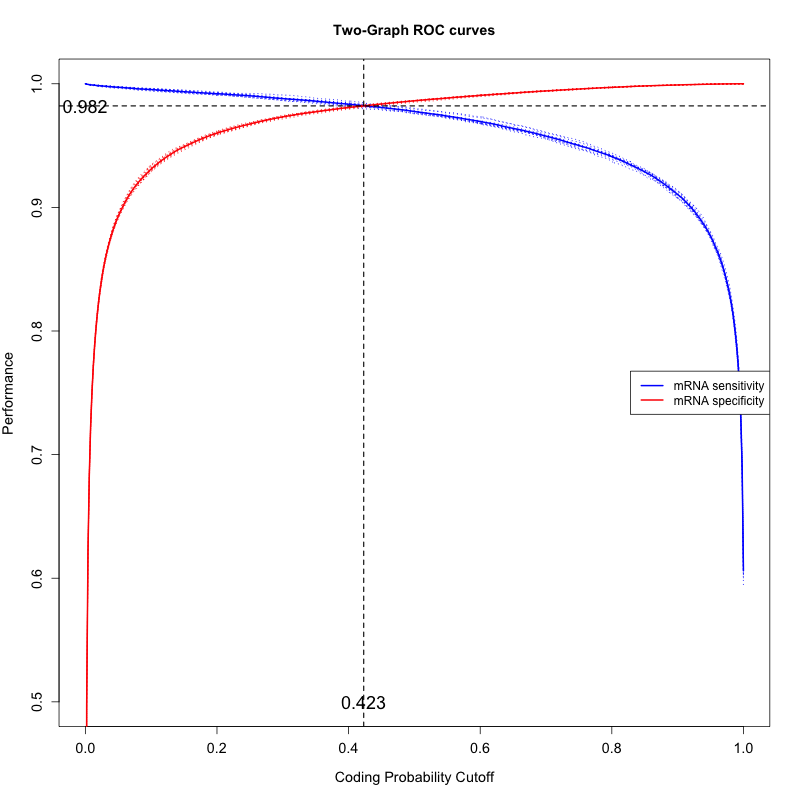

**Figure S1.** Two ROC curves for automatic detection of optimized CPS threshold and user specificity threshold. Red line indicates mRNA specificity. Blue line indicates lncRNA specificity.

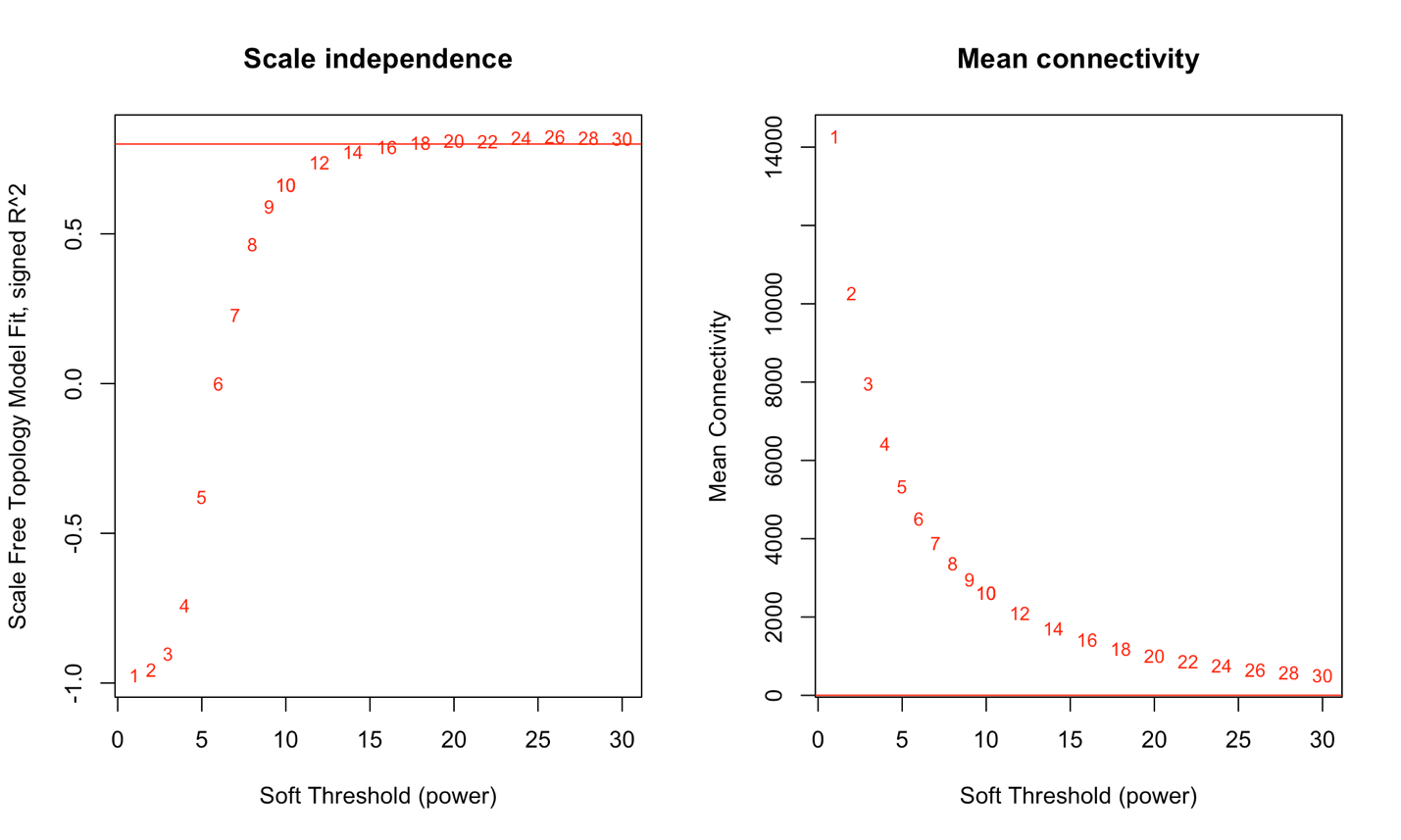

**Figure S2.** Soft threshold filtering. Scale-independence and mean connectivity of the network in different soft-threshold powers. The left panel displays the correlation of soft threshold with scale-free fit index. The right panel displays the influence of soft-threshold power on mean connectivity.

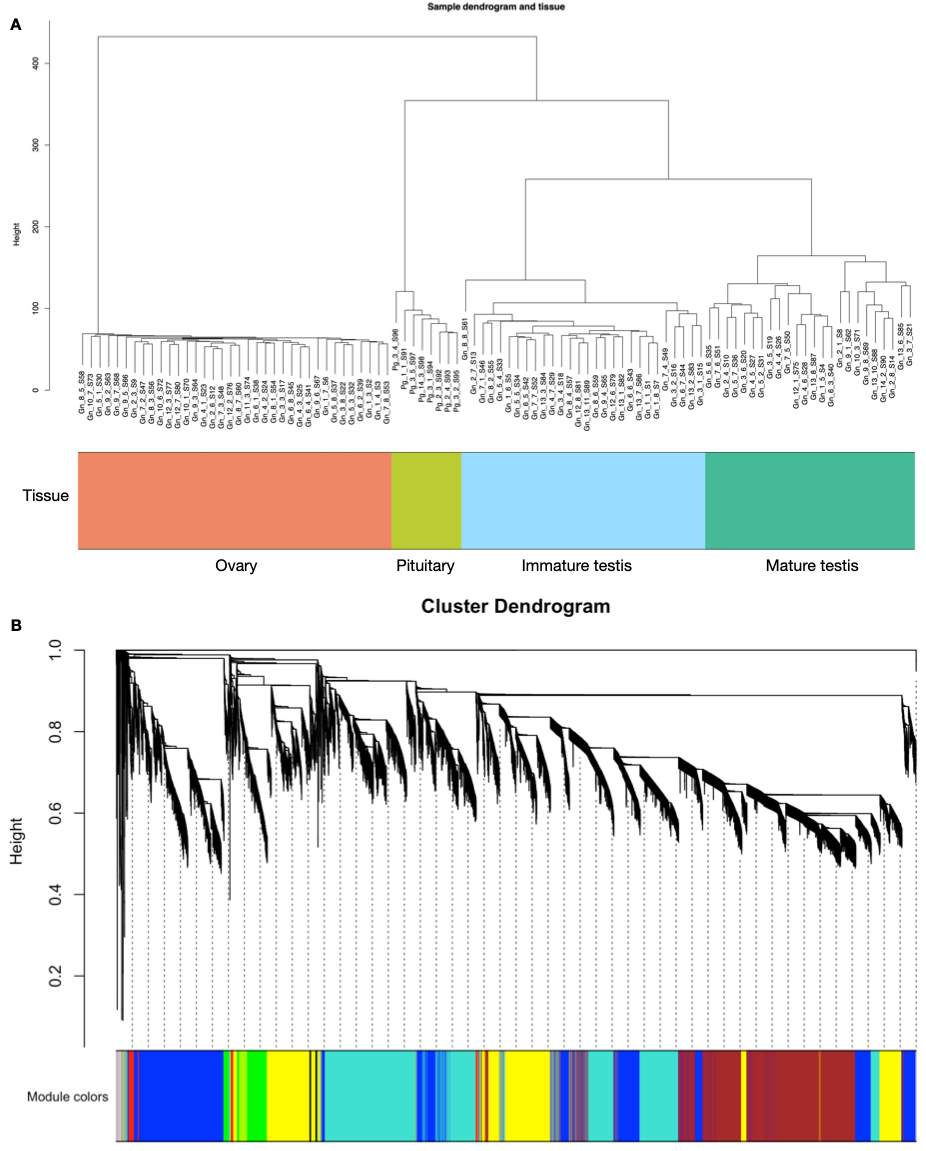

**Figure S3.** WGCNA network construction (A), Sample dendrogram and tissue annotation (B), Cluster dendrogram

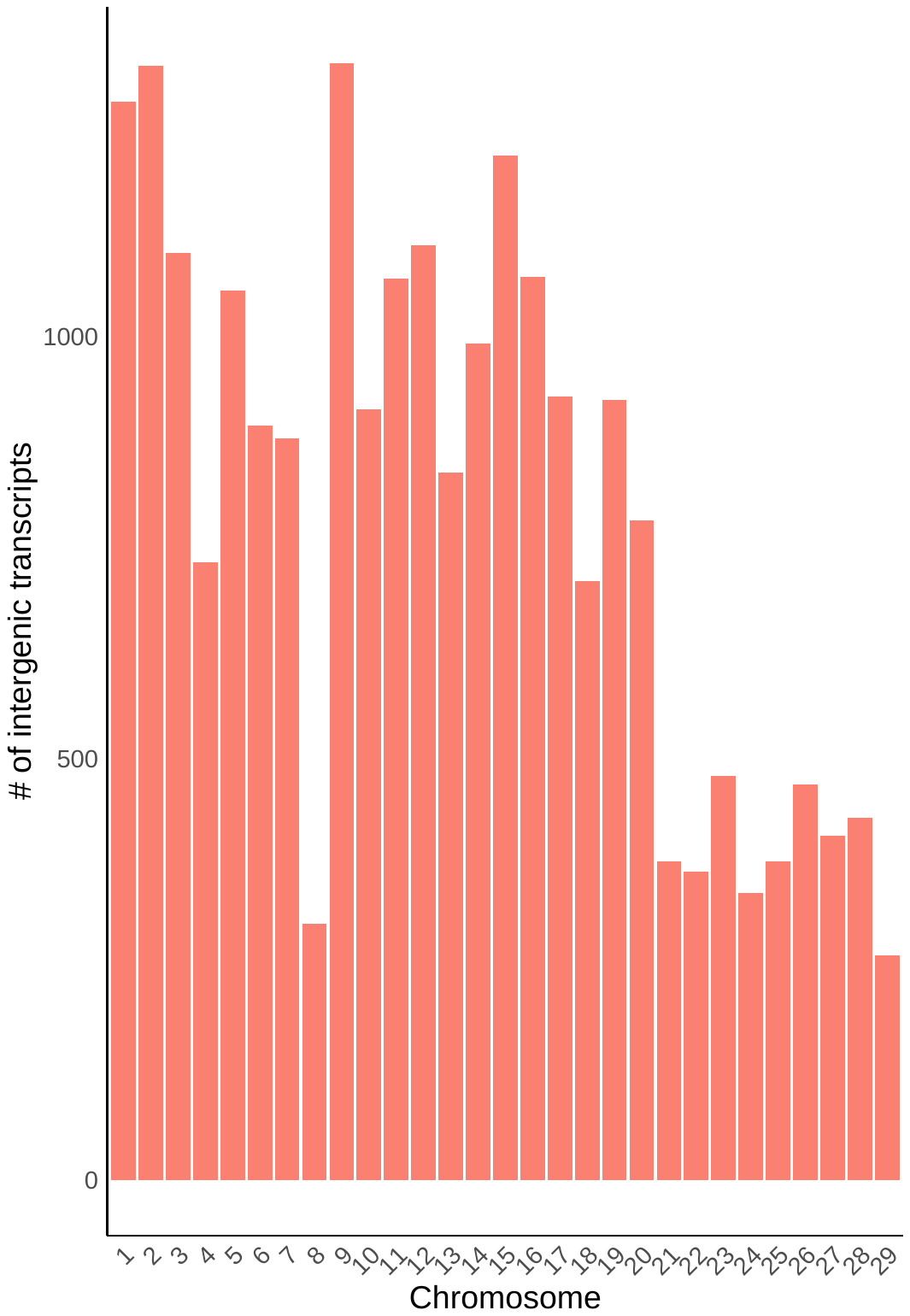

**Figure S4.** Chromosomal distribution of intergenic transcripts

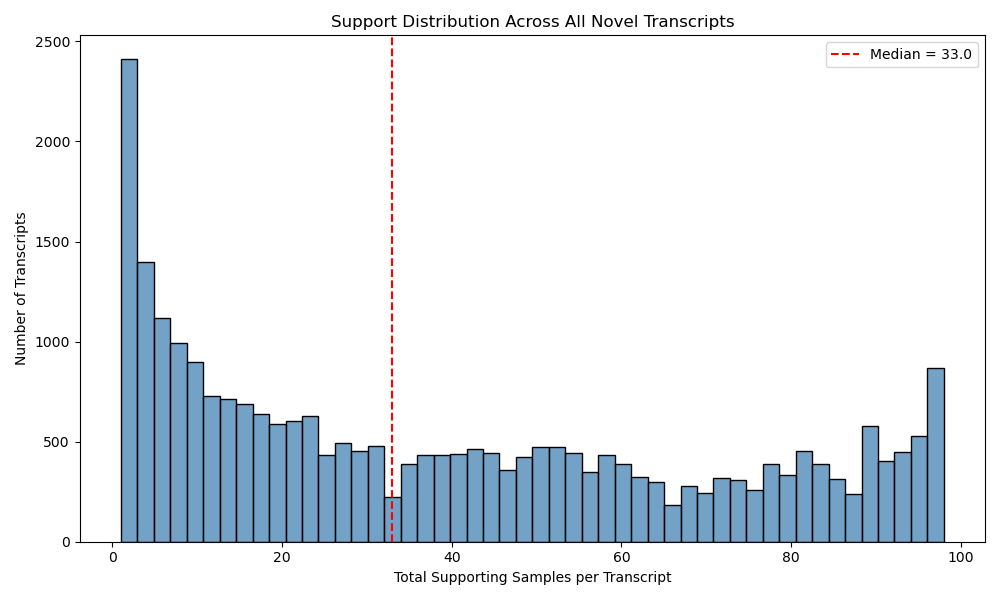

**Figure S5**. Histogram of number of samples supporting intergenic transcripts

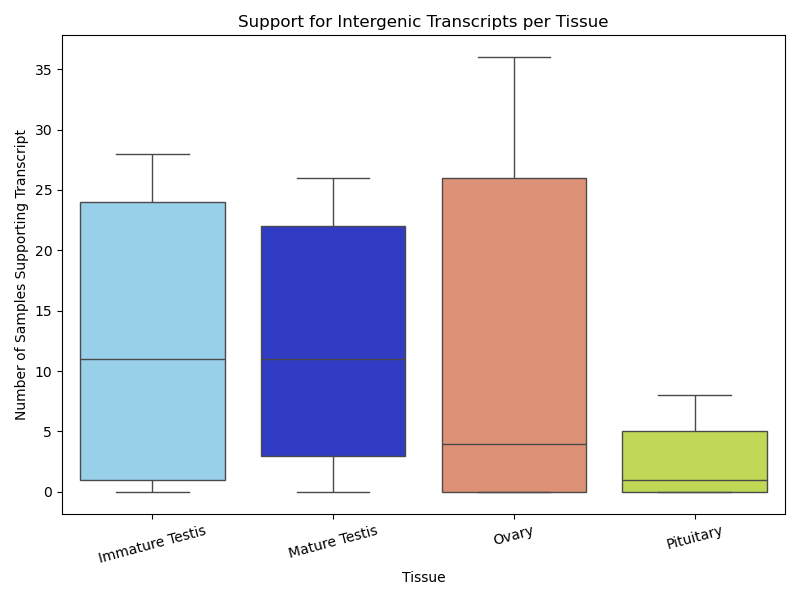

**Figure S6.** Boxplot of the number of samples supporting intergenic transcript per tissue type

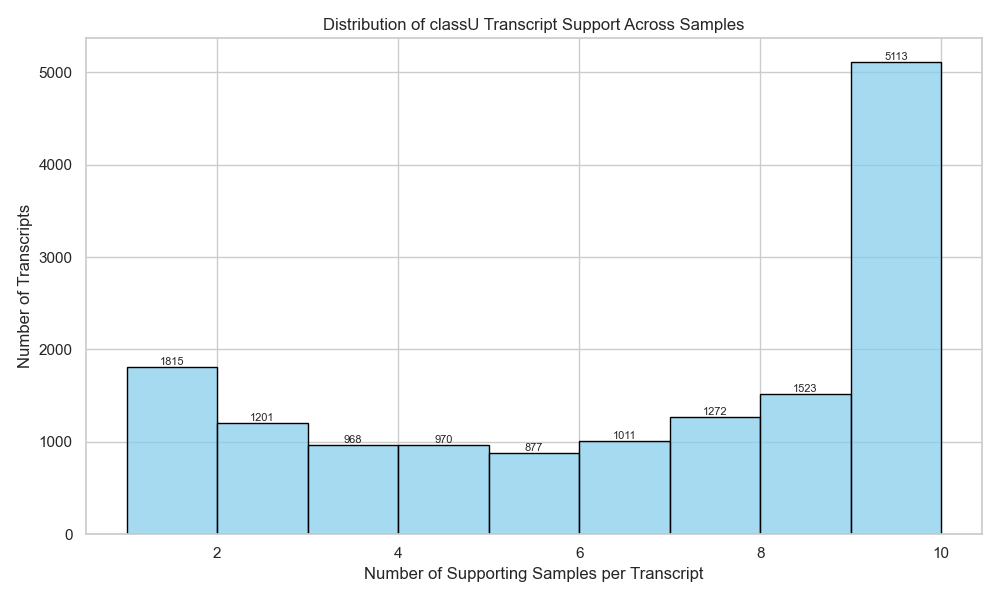

**Figure S7.** Histogram of number of samples supporting intergenic transcripts in PRJNA380580

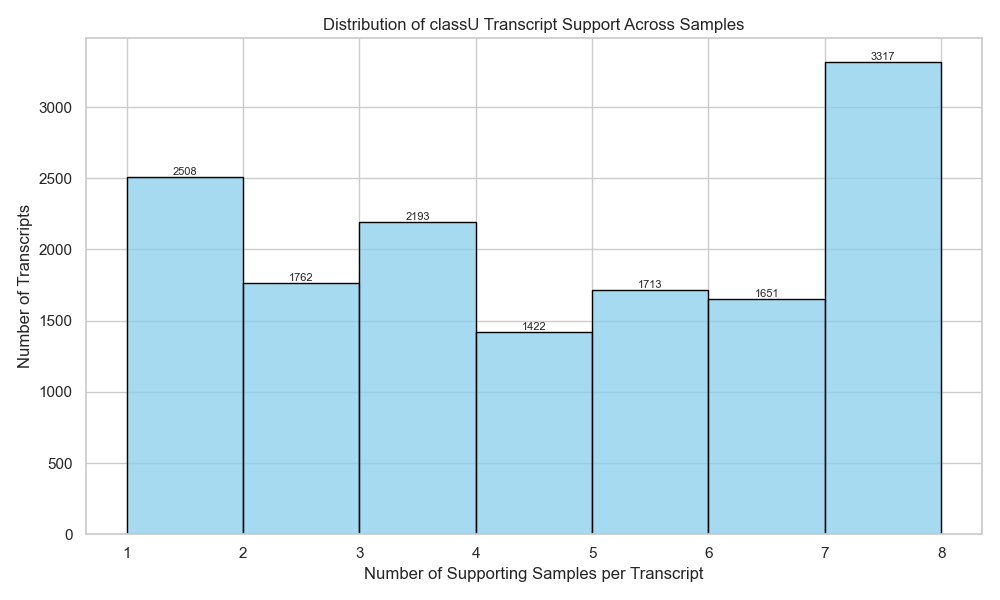

**Figure S8.** Histogram of number of support sample for the intergenic transcripts in PRJNA550414

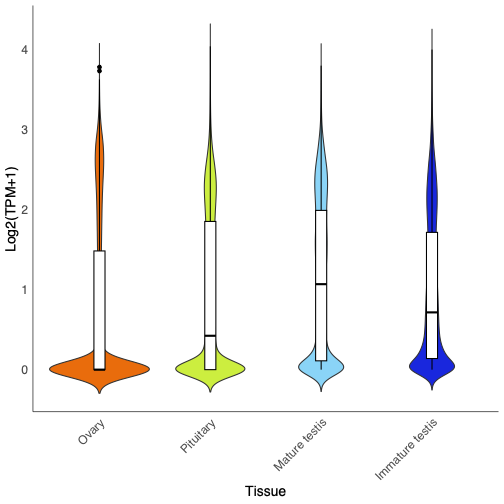

**Figure S9.** A violin plot of RNA expression levels in transcripts per million (TPM) in four Atlantic salmon tissues

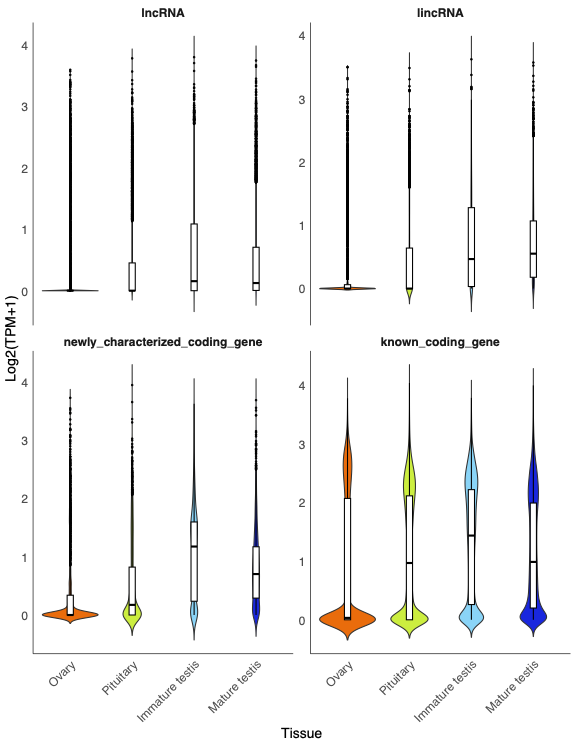

**Figure S10.** Violin plots of known/newly characterized protein-coding gene/lncRNA/lincRNA expression levels in transcripts per million (TPM) across four Atlantic salmon tissues

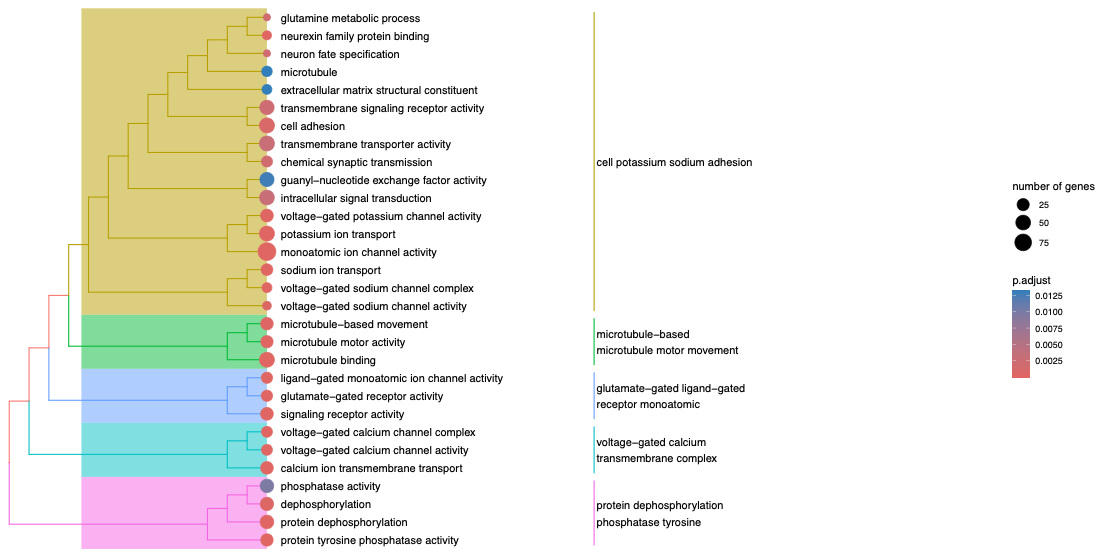

**Figure S11.** Tree plots of the top 30 significantly enriched GO terms in the GO enrichment analysis of yellow module.
