## Supplemental figure and table for "A Comprehensive Analysis of Atlantic Salmon Gonad and Pituitary Transcriptomes Identifies Novel Players in Sexual Maturation": suppelementary_file3_candidate_maturation_related_newly_characterized_gene.docx

**Mturation related candidate newly characterized genes.**

𝜏 value of each loci showing its tissue specificity * indicates loci upregulated in a single tissue, it: immature testis, mt: mature testis, ov: ovary, pt: pituitary

| Module | Geneid | Location | 𝜏 | Tissue | Gene symbol | GO pathway | Reference |
| --- | --- | --- | --- | --- | --- | --- | --- |
| turquoise | MSTRG.19353 | 14:29452500-29454393 | 0.97 | it | *foxq1* | sequence-specific DNA binding | [1] |
| turquoise | MSTRG.36976 | 2:45587769-45591283 | 0.98 | it | *pou3f2* | sequence-specific DNA binding | [2] |
| turquoise | MSTRG.43887 | 22:26980917-26999946 | 0.78 | it | *map4k1* | intracellular signal transduction | [3, 4] |
| turquoise | MSTRG.47909 | 24:42890640-42899662 | 0.93 | it | *chrna2* | transmembrane signaling receptor activity;transmembrane transporter activity | [5] |
| turquoise | MSTRG.50203 | 26:22274896-22279093 | 0.94 | it | *mc1r* | intracellular signal transduction;transmembrane signaling receptor activity | [6, 7] |
| turquoise | MSTRG.51093 | 26:52745846-52752194 | 0.74 | it | *ttn* | intracellular signal transduction;protein tyrosine kinase activity;actin filament binding;myosin complex | [8] |
| turquoise | MSTRG.66318 | 6:43972990-43975586 | 0.75 | it | *hoxb5* | sequence-specific DNA binding | [9] |
| turquoise | MSTRG.71253 | 9:17006614-17024991 | 0.82 | it | *gjb3* | transmembrane transporter activity | [10] |
| turquoise | MSTRG.7678 | 10:101440111-101451026 | 0.89 | it | *nfic* | sequence-specific DNA binding | [11] |
| turquoise | MSTRG.77909 | CAJNNT020001543.1:98775-102564 | 0.76 | it | *map4k1* | intracellular signal transduction | [3, 4] |
| turquoise | MSTRG.9788 | 11:54024351-54027406 | 0.88 | it | *cebpa* | sequence-specific DNA binding | [12] |
| turquoise | MSTRG.77236 | CAJNNT020001308.1:163721-195053 | 0.39 | pt* | *wnk1* | intracellular signal transduction | [13] |
| turquoise | MSTRG.30515 | 17:84363103-84405843 | 0.40 | pt* | *wnk1* | intracellular signal transduction |  |
| blue | MSTRG.14389 | 12:78943353-78949043 | 0.94 | mt | *nfic* | sequence-specific DNA binding | [11] |
| blue | MSTRG.31161 | 18:17331978-17332733 | 0.89 | mt | *nfic* | sequence-specific DNA binding |  |
| blue | MSTRG.41134 | 20:73029699-73038125 | 1.00 | mt | *nfic* | sequence-specific DNA binding |  |
| blue | MSTRG.17899 | 13:96185476-96240120 | 0.48 | mt | *clip1* | cytoskeleton;cytoskeleton organization;actin binding;cell projection;plasma membrane bounded cell projection;neuron projection;microtubule binding;intracellular protein transport | [14] |
| blue | MSTRG.23444 | 15:53950937-53955657 | 1.00 | mt* | *exoc3* | cell projection;plasma membrane bounded cell projection | [15, 16] |
| blue | MSTRG.23516 | 15:58049402-58059974 | 1.00 | mt* | *phc1* | regulation of cell cycle process;cellular response to cytokine stimulus | [17] |
| blue | MSTRG.30638 | 17:86893690-86896024 | 1.00 | mt | *phc1* | regulation of cell cycle process;cellular response to cytokine stimulus |  |
| blue | MSTRG.25955 | 16:28008461-28014267 | 1.00 | mt* | *pop-1* | kinase binding;sequence-specific DNA binding | [18] |
| blue | MSTRG.33633 | 19:29102282-29104208 | 0.55 | pt | *c12orf55 (CFAP54)* | cilium movement involved in cell motility;cilium assembly;cytoskeleton;axoneme;cell projection assembly;cilium;cell projection;plasma membrane bounded cell projection;plasma membrane bounded cell projection assembly;cilium organization;microtubule-based movement | [19] |
| blue | MSTRG.38636 | 2:86521879-86534355 | 1.00 | mt* | *haus7* | cilium assembly;cytoskeleton;cytoskeleton organization;cell projection assembly;plasma membrane bounded cell projection assembly;regulation of cell cycle process;cilium organization | [20] |
| blue | MSTRG.38830 | 2:88262406-88263034 | 1.00 | mt* | *hmgb1* | lipid binding;cytoskeleton organization;cell projection;plasma membrane bounded cell projection;positive regulation of chemotaxis;apoptotic process;membrane raft;neuron projection;GO:0051272;sequence-specific DNA binding;cellular response to cytokine stimulus | [21, 22] |
| blue | MSTRG.43694 | 22:19570003-19570939 | 1.00 | mt | *odc1* | carboxy-lyase activity;carbon-carbon lyase activity | [23, 24] |
| blue | MSTRG.48559 | 25:17958106-17963460 | 0.95 | mt | *arl4c* | cell projection;plasma membrane bounded cell projection | [25] |
| blue | MSTRG.50594 | 26:35236466-35240379 | 1.00 | mt | *dvl2* | transmembrane receptor protein tyrosine kinase signaling pathway;lipid binding;cilium assembly;cytoskeleton;cytoskeleton organization;regulation of actin cytoskeleton organization;axon guidance;cell projection assembly;cilium;kinase binding;cell projection;plasma membrane bounded cell projection;positive regulation of chemotaxis;plasma membrane bounded cell projection assembly;regulation of cell cycle process;cilium organization;GO:0051272 | [26, 27] |
| blue | MSTRG.73428 | 9:92393500-92396027 | 1.00 | mt | *rab11fip5* | cytoskeleton;kinase binding;cellular response to cytokine stimulus | [28] |
| blue | MSTRG.81613 | CAJNNT020003506.1:50387-57638 | 0.52 | mt* | *dnah7* | cilium assembly;cytoskeleton;cytoskeleton organization;axoneme;cell projection assembly;cilium;minus-end-directed microtubule motor activity;dynein intermediate chain binding;dynein light intermediate chain binding;cell projection;plasma membrane bounded cell projection;plasma membrane bounded cell projection assembly;dynein complex;cilium organization;microtubule-based movement | [29] |
| blue | MSTRG.82129 | CAJNNT020004027.1:40133-46756 | 0.51 | mt | *dnah6* | cilium assembly;cytoskeleton;axoneme;cell projection assembly;cilium;minus-end-directed microtubule motor activity;dynein intermediate chain binding;dynein light intermediate chain binding;cell projection;plasma membrane bounded cell projection;plasma membrane bounded cell projection assembly;dynein complex;cilium organization;microtubule-based movement | [30] |
| blue | MSTRG.9377 | 11:35108607-35112465 | 1.00 | mt | *dvl2* | transmembrane receptor protein tyrosine kinase signaling pathway;lipid binding;cilium assembly;cytoskeleton;cytoskeleton organization;regulation of actin cytoskeleton organization;axon guidance;cell projection assembly;cilium;kinase binding;cell projection;plasma membrane bounded cell projection;positive regulation of chemotaxis;plasma membrane bounded cell projection assembly;regulation of cell cycle process;cilium organization;GO:0051272 | [27] |
| brown | MSTRG.26507 | 16:45185343-45187305 | 0.52 | ov* | *ier5* | regulation of transcription by RNA polymerase II | [31] |
| brown | MSTRG.39497 | 20:20812801-20814199 | 0.52 | ov* | *anapc5* | ubiquitin-protein transferase activity | [32] |
| green | MSTRG.33275 | 19:9749110-9754623 | 1.00 | pt | *amer2* | phospholipid binding | [33] |
| green | MSTRG.64355 | 5:85319395-85322944 | 1.00 | pt* | *nbeal2* | secretory granule | [34] |
| green | MSTRG.66383 | 6:45618184-45621858 | 1.00 | pt* | *gh1* | hormone activity;secretory granule;sodium ion transport | [35] |
| green | MSTRG.81880 | CAJNNT020003707.1:118842-162475 | 0.99 | pt | *ugt8* | UDP-glycosyltransferase activity | [36] |
| yellow | MSTRG.49132 | 25:39104782-39107074 | 0.82 | pt | *slc3a2* | calcium ion transmembrane transport;sodium ion transport;transmembrane transporter activity | [37] |
| yellow | MSTRG.51044 | 26:51076808-51080606 | 1.00 | pt | *pten* | protein tyrosine phosphatase activity;protein dephosphorylation;dephosphorylation;chemical synaptic transmission;intracellular signal transduction;phosphatase activity;actin binding;transmembrane receptor protein tyrosine kinase signaling pathway;postsynaptic membrane | [38] |
| yellow | MSTRG.57095 | 3:58358084-58361260 | 0.87 | pt | *krt12* | signaling receptor activity;transmembrane signaling receptor activity;protein dimerization activity | [39] |
| yellow | MSTRG.67178 | 6:72434470-72437104 | 0.91 | pt | *gpr6* | signaling receptor activity;transmembrane signaling receptor activity;intracellular signal transduction | [40] |
| yellow | MSTRG.79860 | CAJNNT020002330.1:2300-8334 | 0.88 | pt | *cpe* | neurexin family protein binding;chemical synaptic transmission | [41] |
