## Supplemental figure and table for "A Comprehensive Analysis of Atlantic Salmon Gonad and Pituitary Transcriptomes Identifies Novel Players in Sexual Maturation": supplementary_file1_scripts.sh

```
#!/bin/bash

######--- 1. fastp v0.23.2: quality control ---######

WD=/ssal_gnpt_rnaseq/read/clean_reads
SD=/ssal_gnpt_rnaseq/read/rawreads

cd $WD;

declare -a samples_all=("Gn_10_1_S70" "Gn_10_3_S71" "Gn_10_6_S72" "Gn_10_7_S73" "Gn_11_3_S74" "Gn_12_1_S75" "Gn_12_2_S76" "Gn_12_3_S77" "Gn_12_5_S78" "Gn_12_6_S79" "Gn_12_7_S80" "Gn_12_8_S81" "Gn_13_10_S88" "Gn_13_11_S89" "Gn_13_1_S82" "Gn_13_2_S83" "Gn_13_3_S84" "Gn_13_6_S85" "Gn_13_7_S86" "Gn_13_8_S87" "Gn_14_2_S90" "Gn_1_1_S1" "Gn_1_3_S2" "Gn_1_4_S3" "Gn_1_5_S4" "Gn_1_6_S5" "Gn_1_7_S6" "Gn_1_8_S7" "Gn_2_1_S8" "Gn_2_3_S9" "Gn_2_4_S10" "Gn_2_5_S11" "Gn_2_6_S12" "Gn_2_7_S13" "Gn_2_8_S14" "Gn_3_1_S15" "Gn_3_2_S16" "Gn_3_3_S17" "Gn_3_4_S18" "Gn_3_5_S19" "Gn_3_6_S20" "Gn_3_7_S21" "Gn_3_8_S22" "Gn_4_1_S23" "Gn_4_2_S24" "Gn_4_3_S25" "Gn_4_4_S26" "Gn_4_5_S27" "Gn_4_6_S28" "Gn_4_7_S29" "Gn_5_1_S30" "Gn_5_2_S31" "Gn_5_3_S32" "Gn_5_4_S33" "Gn_5_5_S34" "Gn_5_6_S35" "Gn_5_7_S36" "Gn_5_8_S37" "Gn_6_1_S38" "Gn_6_2_S39" "Gn_6_3_S40" "Gn_6_4_S41" "Gn_6_5_S42" "Gn_6_6_S43" "Gn_6_7_S44" "Gn_6_8_S45" "Gn_7_1_S46" "Gn_7_2_S47" "Gn_7_3_S48" "Gn_7_4_S49" "Gn_7_5_S50" "Gn_7_6_S51" "Gn_7_7_S52" "Gn_7_8_S53" "Gn_8_1_S54" "Gn_8_2_S55" "Gn_8_3_S56" "Gn_8_4_S57" "Gn_8_5_S58" "Gn_8_6_S59" "Gn_8_7_S60" "Gn_8_8_S61" "Gn_9_1_S62" "Gn_9_2_S63" "Gn_9_3_S64" "Gn_9_4_S65" "Gn_9_5_S66" "Gn_9_6_S67" "Gn_9_7_S68" "Gn_9_8_S69" "Pg_1-1_S91" "Pg_14-3_S98" "Pg_2-3_S92" "Pg_2-4_S93" "Pg_3-1_S94" "Pg_3-2_S95" "Pg_3-4_S96" "Pg_3-5_S97")

for sample in "${samples_all[@]}"
do

fastp --in1 $SD/${sample}.R1.fastq.gz --in2 $SD/${sample}.R2.fastq.gz --out1 $WD/${sample}_clean.R1.fastq.gz --out2 $WD/${sample}_clean.R2.fastq.gz --json=$WD/${sample}_fastp.json --html=$WD/${sample}_fastp.html --compression=6 --thread=5

done

######--- 2.1 STAR v2.7.11a: index for reference genome ---#######

WD=/ssal_gnpt_rnaseq/ref/STAR_Genome_Index
SD=/ssal_gnpt_rnaseq/ref/Ssal-v3.1
GENOME_FASTA=/ssal_gnpt_rnaseq/ref/Salmo_salar-GCA_905237065.2-unmasked.fa
ANNOTATION_GTF=$SD/Salmo_salar-GCA_905237065.2-2021_07-genes.gtf

STAR --runThreadN 4 --runMode genomeGenerate --genomeDir $WD --genomeFastaFiles $GENOME_FASTA --sjdbGTFfile $ANNOTATION_GTF --sjdbOverhang 100 --genomeChrBinNbits 18

######--- 2.2 STAR v2.7.11a: 1st algnment ---#######

WD=/ssal_gnpt_rnaseq/analysis/01mapping/star_1st
SD=/ssal_gnpt_rnaseq/read/clean_reads

INDEX_DIR=/ssal_gnpt_rnaseq/ref/STAR_Genome_Index
ANNOTATION_GTF=/ssal_gnpt_rnaseq/ref/Salmo_salar-GCA_905237065.2-2021_07-genes.gtf

mkdir -p $WD

cd $WD;
declare -a samples_all=("Gn_10_1_S70" "Gn_10_3_S71" "Gn_10_6_S72" "Gn_10_7_S73" "Gn_11_3_S74" "Gn_12_1_S75" "Gn_12_2_S76" "Gn_12_3_S77" "Gn_12_5_S78" "Gn_12_6_S79" "Gn_12_7_S80" "Gn_12_8_S81" "Gn_13_10_S88" "Gn_13_11_S89" "Gn_13_1_S82" "Gn_13_2_S83" "Gn_13_3_S84" "Gn_13_6_S85" "Gn_13_7_S86" "Gn_13_8_S87" "Gn_14_2_S90" "Gn_1_1_S1" "Gn_1_3_S2" "Gn_1_4_S3" "Gn_1_5_S4" "Gn_1_6_S5" "Gn_1_7_S6" "Gn_1_8_S7" "Gn_2_1_S8" "Gn_2_3_S9" "Gn_2_4_S10" "Gn_2_5_S11" "Gn_2_6_S12" "Gn_2_7_S13" "Gn_2_8_S14" "Gn_3_1_S15" "Gn_3_2_S16" "Gn_3_3_S17" "Gn_3_4_S18" "Gn_3_5_S19" "Gn_3_6_S20" "Gn_3_7_S21" "Gn_3_8_S22" "Gn_4_1_S23" "Gn_4_2_S24" "Gn_4_3_S25" "Gn_4_4_S26" "Gn_4_5_S27" "Gn_4_6_S28" "Gn_4_7_S29" "Gn_5_1_S30" "Gn_5_2_S31" "Gn_5_3_S32" "Gn_5_4_S33" "Gn_5_5_S34" "Gn_5_6_S35" "Gn_5_7_S36" "Gn_5_8_S37" "Gn_6_1_S38" "Gn_6_2_S39" "Gn_6_3_S40" "Gn_6_4_S41" "Gn_6_5_S42" "Gn_6_6_S43" "Gn_6_7_S44" "Gn_6_8_S45" "Gn_7_1_S46" "Gn_7_2_S47" "Gn_7_3_S48" "Gn_7_4_S49" "Gn_7_5_S50" "Gn_7_6_S51" "Gn_7_7_S52" "Gn_7_8_S53" "Gn_8_1_S54" "Gn_8_2_S55" "Gn_8_3_S56" "Gn_8_4_S57" "Gn_8_5_S58" "Gn_8_6_S59" "Gn_8_7_S60" "Gn_8_8_S61" "Gn_9_1_S62" "Gn_9_2_S63" "Gn_9_3_S64" "Gn_9_4_S65" "Gn_9_5_S66" "Gn_9_6_S67" "Gn_9_7_S68" "Gn_9_8_S69" "Pg_1-1_S91" "Pg_14-3_S98" "Pg_2-3_S92" "Pg_2-4_S93" "Pg_3-1_S94" "Pg_3-2_S95" "Pg_3-4_S96" "Pg_3-5_S97")

for sample in "${samples_all[@]}"
do

STAR --runMode alignReads --runThreadN 24 --genomeDir $INDEX_DIR —-outFilterIntronMotifs RemoveNoncanonicalUnannotated --readFilesIn $SD/${sample}_clean.R1.fastq.gz $SD/${sample}_clean.R2.fastq.gz --readFilesCommand zcat --sjdbGTFfile $ANNOTATION_GTF --outFileNamePrefix $WD/${sample} --twopassMode None --quantMode GeneCounts --alignIntronMin 20 --alignIntronMax 1000000 --alignSJDBoverhangMin 1 --alignMatesGapMax 1000000 --alignEndsProtrude 10 ConcordantPair --outFilterType BySJout --chimSegmentMin 10 --outSAMtype BAM SortedByCoordinate

done

######--- 2.3 STAR v2.7.11a: reIndex with SJ ---#######

WD=/ssal_gnpt_rnaseq/analysis/01mapping/star_1st/SJ_out_tab
SD=/ssal_gnpt_rnaseq/read/clean_reads

GENOME_FASTA=/ssal_gnpt_rnaseq/ref/Ssal-v3.1/Salmo_salar-GCA_905237065.2-unmasked.fa
ANNOTATION_GTF=/ssal_gnpt_rnaseq/ref/Ssal-v3.1/Salmo_salar-GCA_905237065.2-2021_07-genes.gtf
STA
# Concatenate SJ.out.tab
cat $WD/*SJ.out.tab > $WD/concatenate_SJ.out.tab

SJ_OUT_FILE=$WD/concatenate_SJ.out.tab

# Regenerate index for genome with concatenate SJ.out
GENOME_INDEX_DIR_STAR=/ssal_gnpt_rnaseq/ref/STAR_Genome_Index_with_SJ
mkdir -p ${GENOME_INDEX_DIR_STAR}
cd ${GENOME_INDEX_DIR_STAR}

STAR --runThreadN 20 --runMode genomeGenerate --genomeDir ${GENOME_INDEX_DIR_STAR} --genomeFastaFiles ${GENOME_FASTA} --sjdbGTFfile ${ANNOTATION_GTF} --sjdbFileChrStartEnd ${SJ_OUT_FILE} --limitSjdbInsertNsj 6000000 --sjdbOverhang 100 --genomeChrBinNbits 18

######--- 2.4 STAR v2.7.11a: 2nd alignment ---#######

GENOME_INDEX_DIR_STAR=/ssal_gnpt_rnaseq/ref/STAR_Genome_Index_with_Gn_SJ
GENOME_FASTA=/ssal_gnpt_rnaseq/ref/Ssal-v3.1/Salmo_salar-GCA_905237065.2-unmasked.fa
ANNOTATION_GTF=/ssal_gnpt_rnaseq/ref/Ssal-v3.1/Salmo_salar-GCA_905237065.2-2021_07-genes.gtf
SJ_OUT_FILE=/ssal_gnpt_rnaseq/analysis/01mapping/star_1st/SJ_out_tab/concatenate_SJ.out.tab


WD=/ssal_gnpt_rnaseq/analysis/01mapping/star_2nd
SD=/ssal_gnpt_rnaseq/read/clean_reads

cd $WD;
declare -a samples_all=("Gn_10_1_S70" "Gn_10_3_S71" "Gn_10_6_S72" "Gn_10_7_S73" "Gn_11_3_S74" "Gn_12_1_S75" "Gn_12_2_S76" "Gn_12_3_S77" "Gn_12_5_S78" "Gn_12_6_S79" "Gn_12_7_S80" "Gn_12_8_S81" "Gn_13_10_S88" "Gn_13_11_S89" "Gn_13_1_S82" "Gn_13_2_S83" "Gn_13_3_S84" "Gn_13_6_S85" "Gn_13_7_S86" "Gn_13_8_S87" "Gn_14_2_S90" "Gn_1_1_S1" "Gn_1_3_S2" "Gn_1_4_S3" "Gn_1_5_S4" "Gn_1_6_S5" "Gn_1_7_S6" "Gn_1_8_S7" "Gn_2_1_S8" "Gn_2_3_S9" "Gn_2_4_S10" "Gn_2_5_S11" "Gn_2_6_S12" "Gn_2_7_S13" "Gn_2_8_S14" "Gn_3_1_S15" "Gn_3_2_S16" "Gn_3_3_S17" "Gn_3_4_S18" "Gn_3_5_S19" "Gn_3_6_S20" "Gn_3_7_S21" "Gn_3_8_S22" "Gn_4_1_S23" "Gn_4_2_S24" "Gn_4_3_S25" "Gn_4_4_S26" "Gn_4_5_S27" "Gn_4_6_S28" "Gn_4_7_S29" "Gn_5_1_S30" "Gn_5_2_S31" "Gn_5_3_S32" "Gn_5_4_S33" "Gn_5_5_S34" "Gn_5_6_S35" "Gn_5_7_S36" "Gn_5_8_S37" "Gn_6_1_S38" "Gn_6_2_S39" "Gn_6_3_S40" "Gn_6_4_S41" "Gn_6_5_S42" "Gn_6_6_S43" "Gn_6_7_S44" "Gn_6_8_S45" "Gn_7_1_S46" "Gn_7_2_S47" "Gn_7_3_S48" "Gn_7_4_S49" "Gn_7_5_S50" "Gn_7_6_S51" "Gn_7_7_S52" "Gn_7_8_S53" "Gn_8_1_S54" "Gn_8_2_S55" "Gn_8_3_S56" "Gn_8_4_S57" "Gn_8_5_S58" "Gn_8_6_S59" "Gn_8_7_S60" "Gn_8_8_S61" "Gn_9_1_S62" "Gn_9_2_S63" "Gn_9_3_S64" "Gn_9_4_S65" "Gn_9_5_S66" "Gn_9_6_S67" "Gn_9_7_S68" "Gn_9_8_S69" "Pg_1-1_S91" "Pg_14-3_S98" "Pg_2-3_S92" "Pg_2-4_S93" "Pg_3-1_S94" "Pg_3-2_S95" "Pg_3-4_S96" "Pg_3-5_S97")

for sample in "${samples_all[@]}"
do

STAR --runMode alignReads --runThreadN 24 --genomeDir $GENOME_INDEX_DIR_STAR --outFilterIntronMotifs RemoveNoncanonicalUnannotated --readFilesIn $SD/${sample}_clean.R1.fastq.gz $SD/${sample}_clean.R2.fastq.gz --readFilesCommand zcat --sjdbGTFfile $ANNOTATION_GTF --outFileNamePrefix $WD/${sample} --twopassMode None --quantMode GeneCounts --alignIntronMin 20 --alignIntronMax 1000000 --alignSJDBoverhangMin 1 --alignMatesGapMax 1000000 --alignEndsProtrude 10 ConcordantPair --outFilterType BySJout --chimSegmentMin 10 --outSAMtype BAM SortedByCoordinate --limitSjdbInsertNsj 6000000 --sjdbFileChrStartEnd $SJ_OUT_FILE


########---- 3. StringTie v2.2.1 : transcriptome assembly ----##################

WD=/ssal_gnpt_rnaseq/analysis/02stringtie_Ssal_gnpt
SD=/ssal_gnpt_rnaseq/analysis/01mapping/star_2nd
ANNOTATION_GTF=/ssal_gnpt_rnaseq/ref/Ssal-v3.1/Salmo_salar-GCA_905237065.2-2021_07-genes.gtf

cd $WD;

declare -a samples_all=("Gn_10_1_S70" "Gn_10_3_S71" "Gn_10_6_S72" "Gn_10_7_S73" "Gn_11_3_S74" "Gn_12_1_S75" "Gn_12_2_S76" "Gn_12_3_S77" "Gn_12_5_S78" "Gn_12_6_S79" "Gn_12_7_S80" "Gn_12_8_S81" "Gn_13_10_S88" "Gn_13_11_S89" "Gn_13_1_S82" "Gn_13_2_S83" "Gn_13_3_S84" "Gn_13_6_S85" "Gn_13_7_S86" "Gn_13_8_S87" "Gn_14_2_S90" "Gn_1_1_S1" "Gn_1_3_S2" "Gn_1_4_S3" "Gn_1_5_S4" "Gn_1_6_S5" "Gn_1_7_S6" "Gn_1_8_S7" "Gn_2_1_S8" "Gn_2_3_S9" "Gn_2_4_S10" "Gn_2_5_S11" "Gn_2_6_S12" "Gn_2_7_S13" "Gn_2_8_S14" "Gn_3_1_S15" "Gn_3_2_S16" "Gn_3_3_S17" "Gn_3_4_S18" "Gn_3_5_S19" "Gn_3_6_S20" "Gn_3_7_S21" "Gn_3_8_S22" "Gn_4_1_S23" "Gn_4_2_S24" "Gn_4_3_S25" "Gn_4_4_S26" "Gn_4_5_S27" "Gn_4_6_S28" "Gn_4_7_S29" "Gn_5_1_S30" "Gn_5_2_S31" "Gn_5_3_S32" "Gn_5_4_S33" "Gn_5_5_S34" "Gn_5_6_S35" "Gn_5_7_S36" "Gn_5_8_S37" "Gn_6_1_S38" "Gn_6_2_S39" "Gn_6_3_S40" "Gn_6_4_S41" "Gn_6_5_S42" "Gn_6_6_S43" "Gn_6_7_S44" "Gn_6_8_S45" "Gn_7_1_S46" "Gn_7_2_S47" "Gn_7_3_S48" "Gn_7_4_S49" "Gn_7_5_S50" "Gn_7_6_S51" "Gn_7_7_S52" "Gn_7_8_S53" "Gn_8_1_S54" "Gn_8_2_S55" "Gn_8_3_S56" "Gn_8_4_S57" "Gn_8_5_S58" "Gn_8_6_S59" "Gn_8_7_S60" "Gn_8_8_S61" "Gn_9_1_S62" "Gn_9_2_S63" "Gn_9_3_S64" "Gn_9_4_S65" "Gn_9_5_S66" "Gn_9_6_S67" "Gn_9_7_S68" "Gn_9_8_S69" "Pg_1-1_S91" "Pg_14-3_S98" "Pg_2-3_S92" "Pg_2-4_S93" "Pg_3-1_S94" "Pg_3-2_S95" "Pg_3-4_S96" "Pg_3-5_S97")

for sample  in  "${samples_all[@]}"
do
output_file="${sample}_stringtie.gtf"
stringtie $SD/${sample}_Aligned.sortedByCoord.out.bam -p 8 --rf -e -B -o $WD/${sample}_stringtie.gtf -G $ANNOTATION_GTF  -A $WD/${sample}_stringtie.tab -C $WD/${sample}_cov_refs.gtf >> $WD/Stringtie_console;

done

#########--- 4. StringTie v2.2.1 : merge annotations ------###############

  ## the Ensembl annotation file used for read alignment and transcript assembly with STAR and Stringtie, respectively ##

WD=/ssal_gnpt_rnaseq/analysis/02stringtie_Ssal_gnpt
ANNOTATION_GTF=/ssal_gnpt_rnaseq/ref/Ssal-v3.1/Salmo_salar-GCA_905237065.2-2021_07-genes.gtf

cd $WD/

find $WD -name "*.gtf" > $WD/transcripts_to_merge.txt

stringtie -p 8 --merge -G $ANNOTATION_GTF -o $WD/98samples_all_stringtie_merged.gtf $WD/transcripts_to_merge.txt

###########---- 5. gffcompare v0.12.6 : compare with reference annotation ------###########

WD=/ssal_gnpt_rnaseq/analysis/02stringtie_Ssal_gnpt
ANNOTATION_GTF=/ssal_gnpt_rnaseq/ref/Ssal-v3.1/Salmo_salar-GCA_905237065.2-2021_07-genes.gtf


cd $WD;

gffcompare -r $ANNOTATION_GTF -o $WD/98samples_all_stringtie_merged.gtf $WD/98samples_all_stringtie_merged.gtf.annotated.gtf

######################################################################################################################

###########---- 6. gffread v0.12.6 : compare with reference annotation ------###########

WD=/ssal_gnpt_rnaseq/analysis/02stringtie_Ssal_gnpt/classU
SD=/ssal_gnpt_rnaseq/analysis/02stringtie_Ssal_gnpt

ANNOTATION_GTF=/ssal_gnpt_rnaseq/ref/Salmo_salar-GCA_905237065.2-2021_07-genes.gtf
GENOME_FASTA=/ssal_gnpt_rnaseq/ref/Salmo_salar-GCA_905237065.2-unmasked.fa
gffcompareTMAP=$SD/98samples_all_stringtie_merged.gtf.annotated.gtf.tmap
gffcompareGTF=$SD/98samples_all_stringtie_merged.gtf.annotated.gtf

classuTMAP=$WD/98samples_all_stringtie_merged.annotated.classU.tmap
classuGTF=$WD/98samples_all_stringtie_merged.annotated.classU.gtf

    ## extract class-u transcripts gtf
cat $gffcompareTMAP | awk '$3 == "u"' > $classuTMAP

~/extract_gtf.sh $classuTMAP $gffcompareGTF $classuGTF

gffread $classuGTF -g $GENOME_FASTA -w $WD/98samples_all_stringtie_merged.annotated.classU.fasta

###########---- 7. TransDecoder v5.7.1 : predict coding regions within transcripts ------###########

WD=/ssal_gnpt_rnaseq/analysis/03novel_pc_transcipts/
SD=/ssal_gnpt_rnaseq/analysis/02stringtie_Ssal_gnpt/classU
classuFASTA=$SD/98samples_all_stringtie_merged.annotated.classU.fasta
longestORF=$WD/98samples_all_stringtie_merged.annotated.classU.fasta.transdecoder_dir/longest_orfs.pep
PFAM_HITS=$WD/pfam.dotblout

    ##1. extract the long open reading frames

TransDecoder.LongOrfs -t $claasuFASTA

    ##2. identify ORFs with homology to known proteins of pfam searches with HMMSCAN

hmmscan --cpu $12 --domtblout pfam.domtbout ./Pfam-A.hmm $longestORF  > common_pfam.log

    ##3. predict the likely coding regions

TransDecoder.Predict -t $claasuFASTA --retain_pfam_hits $PFAM_HITS --single_best_only


###########---- 8. eggNOG-mapper: functional annotation intergenic transcripts ------###########

WD=/ssal_gnpt_rnaseq/analysis/03novel_pc_transcipts/emapper_annotation
SD=/ssal_gnpt_rnaseq/analysis/03novel_pc_transcipts/

EMAPPER_DIR=/usr/local/Caskroom/miniconda/base/envs/eggnog/bin/
EGGNOG_DATA=/usr/local/Caskroom/miniconda/base/lib/python3.10/site-packages/eggnog-mapper-data

mkdir $WD
cd $WD

$EMAPPER_DIR/emapper.py --data_dir $EGGNOG_DATA --cpu 8 -i $SD/98samples_all_stringtie_merged.annotated.classU.fasta.transdecoder.pep -o 98samples_all_stringtie_merged.annotated.classU.fasta.transdecoder._empper --output_dir $WD

######################################################################################################################mm

###########---- 9. FEElnc : lncRNAs analysis ------###########

WD=/ssal_gnpt_rnaseq/analysis/04FEElnc_Ssal_gnpt
gffcompareGTF=/ssal_gnpt_rnaseq/02stringtie_Ssal_gnpt/98samples_all_stringtie_merged.gtf.annotated.gtf
GENOME_FASTA=/ssal_gnpt_rnaseq/ref/Salmo_salar-GCA_905237065.2-unmasked.fa
FEELnc_DIR=/Users/xindhuan/FEElnc_dir/

cd $WD;

    ######## moudle1: extract, filter candidate transcripts #######
$FEELnc_DIR/bin/FEELnc_filter.pl -p 12 -i $gffcompareGTF -a $GENOME_FASTA -b transcript_biotype=protein_coding \
 --monoex=-1 -o $WD/FEELnc_filtered_MERGED_gffcompare.out > $WD/Candidate_lncRNA_FEELnc_Filtered.gtf

    ######## moudle2: compute the coding potential of canadidate transcripts #######

$FEELnc/bin/FEELnc_codpot.pl -p 12 -i $WD/Candidate_lncRNA_FEELnc_Filtered.gtf -a $GTFensembl95 -g $GENOME_FASTA  -b transcript_biotype=protein_coding --outdir $WD/feelnc_codpot_out --outname=Ssal_gnpt_Feelnc --mode=shuffle

   ######## moudle3: classify lncRNAs based on their genomic localization with others transcripts.
 #######

$FEELnc/bin/FEELnc_classifier.pl --biotype -i $WD/feelnc_codpot_out/Ssal_gnpt_Feelnc.lncRNA.gtf -a $GENOME_FASTA > $WD/Ssal_gnpt_lncRNA_classes.txt

###########---- 10. gffcompare v0.12.6 : lincRNA identification ------###########

WD=/ssal_gnpt_rnaseq/analysis/04FEElnc_Ssal_gnpt/gffcompareLncRNA
SD=/ssal_gnpt_rnaseq/analysis/04FEElnc_Ssal_gnpt/feelnc_codpot_out
ANNOTATION_GTF=/ssal_gnpt_rnaseq/ref/Salmo_salar-GCA_905237065.2-2021_07-genes.gtf

gffcompare -r $ANNOTATION_GTF -o $WD/Ssal_gnpt_Feelnc_gffcompare.lncRNA.gtf $SD/Ssal_gnpt_Feelnc.lncRNA.gtf

######################################################################################################################mm

###########---- featureCounts of subread v2.0.6 : counts mapped reads for genes ------###########

WD=/ssal_gnpt_rnaseq/analysis/05quantification
SD=/ssal_gnpt_rnaseq/analysis/01mapping/star_2nd

featureCounts=/Users/xindhuan/subread-2.0.6-macOS-x86_64/bin/featureCounts
gffcompareGTF=/ssal_gnpt_rnaseq/02stringtie_Ssal_gnpt/98samples_all_stringtie_merged.gtf.annotated.gtf

cd $WD;

declare -a samples_all=("Gn_10_1_S70" "Gn_10_3_S71" "Gn_10_6_S72" "Gn_10_7_S73" "Gn_11_3_S74" "Gn_12_1_S75" "Gn_12_2_S76" "Gn_12_3_S77" "Gn_12_5_S78" "Gn_12_6_S79" "Gn_12_7_S80" "Gn_12_8_S81" "Gn_13_10_S88" "Gn_13_11_S89" "Gn_13_1_S82" "Gn_13_2_S83" "Gn_13_3_S84" "Gn_13_6_S85" "Gn_13_7_S86" "Gn_13_8_S87" "Gn_14_2_S90" "Gn_1_1_S1" "Gn_1_3_S2" "Gn_1_4_S3" "Gn_1_5_S4" "Gn_1_6_S5" "Gn_1_7_S6" "Gn_1_8_S7" "Gn_2_1_S8" "Gn_2_3_S9" "Gn_2_4_S10" "Gn_2_5_S11" "Gn_2_6_S12" "Gn_2_7_S13" "Gn_2_8_S14" "Gn_3_1_S15" "Gn_3_2_S16" "Gn_3_3_S17" "Gn_3_4_S18" "Gn_3_5_S19" "Gn_3_6_S20" "Gn_3_7_S21" "Gn_3_8_S22" "Gn_4_1_S23" "Gn_4_2_S24" "Gn_4_3_S25" "Gn_4_4_S26" "Gn_4_5_S27" "Gn_4_6_S28" "Gn_4_7_S29" "Gn_5_1_S30" "Gn_5_2_S31" "Gn_5_3_S32" "Gn_5_4_S33" "Gn_5_5_S34" "Gn_5_6_S35" "Gn_5_7_S36" "Gn_5_8_S37" "Gn_6_1_S38" "Gn_6_2_S39" "Gn_6_3_S40" "Gn_6_4_S41" "Gn_6_5_S42" "Gn_6_6_S43" "Gn_6_7_S44" "Gn_6_8_S45" "Gn_7_1_S46" "Gn_7_2_S47" "Gn_7_3_S48" "Gn_7_4_S49" "Gn_7_5_S50" "Gn_7_6_S51" "Gn_7_7_S52" "Gn_7_8_S53" "Gn_8_1_S54" "Gn_8_2_S55" "Gn_8_3_S56" "Gn_8_4_S57" "Gn_8_5_S58" "Gn_8_6_S59" "Gn_8_7_S60" "Gn_8_8_S61" "Gn_9_1_S62" "Gn_9_2_S63" "Gn_9_3_S64" "Gn_9_4_S65" "Gn_9_5_S66" "Gn_9_6_S67" "Gn_9_7_S68" "Gn_9_8_S69" "Pg_1-1_S91" "Pg_14-3_S98" "Pg_2-3_S92" "Pg_2-4_S93" "Pg_3-1_S94" "Pg_3-2_S95" "Pg_3-4_S96" "Pg_3-5_S97")

for sample  in  "${samples_all[@]}"
do

output_file="${sample}_featureCounts.txt"
$featureCounts -T 8 -p -a $gffcompareGTF -t exon -g gene_id -o $output_file -s 2 --largestOverlap -M --fraction $SD/${sample}_Aligned.sortedByCoord.out.bam >> $WD/summary_console;

done
```
