## Supplemental figure and table for "A Comprehensive Analysis of Atlantic Salmon Gonad and Pituitary Transcriptomes Identifies Novel Players in Sexual Maturation": supplementary_file2_Rscripts.R

```
# load library
library("DESeq2")
library("magrittr") 
library("tidyverse")
library("tibble")
library("ggplot2")
library("pheatmap")
library("UpSetR")
library("plyr")
library("reshape2")
library("PCAtools")
library("dplyr")
library("gplots")
library("ggpubr")
library("grid")
library("gridExtra")
library("genefilter")
library("clusterProfiler") 
library("enrichplot")
library("GO.db")
library("WGCNA")
# setThreads
enableWGCNAThreads()
allowWGCNAThreads()

#####-------1. DEG analysis

## DESeq2: Differentially expressed gene analysis 
### prepare data
ssalCts=("/ssal_gnpt_rnaseq/analysis/05quantification/ssal_gnpt_read_count.txt")
ssalAnno=("/ssal_gnpt_rnaseq/analysis/SsalGnPt_samples.csv")

cts <- as.matrix(read.csv(ssalCts, sep = "\t", row.names = "Geneid", na.strings = "NA"))
coldata <- read.csv(ssalAnno, row.names = 2, na.strings = "NA")

coldata$tissue <- factor(coldata$tissue)

rm(ssalAnno, ssalCts)

#### re-arrange sample order
all(rownames(coldata) == colnames(cts))

all(rownames(coldata) %in% colnames(cts))

cts <- cts[, rownames(coldata)]
all(rownames(coldata) == colnames(cts))
dim(cts)

#### independent filtering: required that the number of counts be more than 3 in more than two of the samples. 
cts <- cts[rowSums(cts>3) > 2,]
dim(cts) 

####add DESeq object
dds <- DESeqDataSetFromMatrix(countData = round(cts),
                              colData = coldata,
                              design = as.formula(~tissue))

table(dds$tissue)
dds

#### estimate size factors
dds <- estimateSizeFactors(dds)

sf_plot <- ggplot(tibble(
  `size factor` = dds$sizeFactor,
  `sum` = colSums(cts)), aes(x = `size factor`, y = `sum`)) +
  geom_point() + 
  theme_minimal() +
  theme(
    panel.grid.major = element_blank(),  # Remove major grid lines
    panel.grid.minor = element_blank(),  # Remove minor grid lines
    panel.border = element_blank(),      # Remove panel border
    axis.line = element_line(size = 0.5, color = "black"),# axis lines
    axis.text = element_text(size = 10), # Adjust axis text size
    axis.title = element_text(size = 12) # Adjust axis title size
  )

print(sf_plot)

#### conduct pca
vst <- assay(vst(dds))
p <- pca(vst, metadata = coldata)
biplot(p, showLoadings = T, colby = 'tissue', legendPosition = 'right', max.overlaps = 50, ntopLoadings = 20, lab = NULL)


# remove outliers
columns_to_keep <- colnames(cts)[!colnames(cts) %in% c("Gn_12_5_S78", "Gn_2_5_S11")]

cts <- cts[, columns_to_keep]

dim(cts)
colnames(cts)

samples_to_keep <- rownames(coldata)[!rownames(coldata) %in% c("Gn_12_5_S78", "Gn_2_5_S11")]

coldata <- coldata[samples_to_keep, ]

dim(coldata)
head(coldata)
rm(columns_to_keep, samples_to_keep)

#### update DESeq object
dds <- DESeqDataSetFromMatrix(countData = round(cts),
                              colData = coldata,
                              design = as.formula(~tissue))

table(dds$tissue)
dds


### run DESeq2 modle
# run model
dds <- DESeq(dds, betaPrior = FALSE)

resultsNames(dds)

#### DE analysis, pairwise comparisons
MtIt <- results(dds, contrast = c("tissue", "mat_ts", "imm_ts"),
                independentFiltering = TRUE, alpha = 0.01, pAdjustMethod = "BH", parallel = TRUE)
summary(MtIt)

MtOv <- results(dds, contrast = c("tissue", "mat_ts", "ov"),
                independentFiltering = TRUE, alpha = 0.01, pAdjustMethod = "BH", parallel = TRUE)
summary(MtOv)

ItOv <- results(dds, contrast = c("tissue", "imm_ts", "ov"),
                independentFiltering = TRUE, alpha = 0.01, pAdjustMethod = "BH", parallel = TRUE)
summary(ItOv)

PtIt <- results(dds, contrast = c("tissue", "pt", "imm_ts"),
                independentFiltering = TRUE, alpha = 0.01, pAdjustMethod = "BH", parallel = TRUE)
summary(PtIt)

PtMt <- results(dds, contrast = c("tissue", "pt", "mat_ts"),
                independentFiltering = TRUE, alpha = 0.01, pAdjustMethod = "BH", parallel = TRUE)
summary(PtMt)

PtOv <- results(dds, contrast = c("tissue", "pt", "ov"),
                independentFiltering = TRUE, alpha = 0.01, pAdjustMethod = "BH", parallel = TRUE)
summary(PtOv)

#####-------2. WGCNA

### normalize counts with DESeq2
vsd <- varianceStabilizingTransformation(dds)

wpn_vsd <- getVarianceStabilizedData(dds)
rv_wpn <- rowVars(wpn_vsd)
summary(rv_wpn)

### input matrix (normalized and 75 quantile expression)
q75_wpn <- quantile( rowVars(wpn_vsd), .75)  # <= 75 quantile to reduce dataset

#keeps only the genes (rows) whose variance exceeds the 75th percentile
expr_normalized <- wpn_vsd[ rv_wpn > q75_wpn, ]
expr_normalized[1:5,1:10]
dim(expr_normalized)

# transpose the data and prepare the dataset for WGCNA
input_mat = t(expr_normalized)

# determine whether there are too many missing values in samples and genes;
gsg<-goodSamplesGenes(input_mat, verbose = 3)
gsg$allOK

### conduct network
#### soft threshold parameters
powers = c(c(1:20), seq(from = 22, to = 30, by = 2))

## unsigned network
sft = pickSoftThreshold(
  input_mat, 
  networkType = "unsigned", 
  powerVector = powers,
  verbose = 5
  )

sft$fitIndices
par(mfrow = c(1,2));  
cex1 = 0.9;  
# plot for "Scale Independence"
plot(sft$fitIndices[, 1],
     -sign(sft$fitIndices[, 3]) * sft$fitIndices[, 2],
     xlab = "Soft Threshold (power)",
     ylab = "Scale Free Topology Model Fit, signed R^2",
     type = "n",
     main = "Scale independence")
text(sft$fitIndices[, 1],
     -sign(sft$fitIndices[, 3]) * sft$fitIndices[, 2],
     labels = powers, cex = cex1, col = "red")
abline(h = 0.8, col = "red")

# plot for "Mean Connectivity"
plot(sft$fitIndices[, 1],
     sft$fitIndices[, 5],
     xlab = "Soft Threshold (power)",
     ylab = "Mean Connectivity",
     type = "n",
     main = "Mean connectivity")
text(sft$fitIndices[, 1],
     sft$fitIndices[, 5],
     labels = powers,
     cex = cex1, col = "red")
abline(h = 0.8, col = "red") 

picked_power = 16

nGenes = ncol(input_mat)

temp_cor <- cor       

cor <- WGCNA::cor

### blockwiseModule
netwk <- blockwiseModules(input_mat,  
                          power = picked_power,            
                          TOMType = "unsigned",
                          deepSplit = 2.5,
                          pamRespectsDendro = F,
                          minModuleSize = 30,
                          maxBlockSize = nGenes,
                          reassignThreshold = 0,
                          mergeCutHeight = 0.25,
                          saveTOMs = T,
                          saveTOMFileBase = "gmssal",
                          numericLabels = T,
                          verbose = 3)

#### check TOP resutls
names(netwk)
table(netwk$colors)

moduleLabels = netwk$colors
moduleColors = labels2colors(netwk$colors)
MEs = netwk$MEs

### correlate module eigengenes with tissue
#### module-tissue by binarizing the tissue variable
# binarize the categorical variavle
binaryTissue <- binarizeCategoricalVariable(coldata$tissue, includeLevelVsAll = TRUE, includePairwise = FALSE );
dim(binaryTissue)

MES0 <- moduleEigengenes(input_mat, moduleColors)$eigengenes
MEs = orderMEs(MES0)

# recalculate the correlations with the binarized variables with cor() Fast calculations of Pearson correlation in WGCNA
moduleTraitCor <- cor(MEs, binaryTissue, use = "p")

moduleTraitPvalue <- corPvalueStudent(moduleTraitCor,nSamples)
textMatrix = paste(signif(moduleTraitCor,2),"\n(",
                     signif(moduleTraitPvalue,1),")",sep = "")


#####-------3. Tissue-specificity analysis

tissueNames <- c("mature_testis", "immature_testis", "ovary", "pituitary")
orgExpression <- read.table("/ssal_gnpt_rnaseq/analysis/featureCount_Ssal_gnpt_TrimmedInput/ssal_gnpt_TPM.txt", header = T, sep = '\t')

## check tpm matrix
dim(orgExpression)

##! we need to remove the outliers "Gn_12_5_S78", "Gn_2_5_S11" which are TS_Mat_10, TS_Mat_13
columns_to_keep <- colnames(orgExpression)[!colnames(orgExpression) %in% c("TS_Mat_10", "TS_Mat_13")]
orgExpression <- orgExpression[, columns_to_keep]
dim(orgExpression)

#Functions# 
##! the functions scripts were based on (Kryuchkova-Mostacci & Robinson-Rechavi, 2017)
fReplicateMean <- function(x) {
  ovary_cols <- grep("^OV_", names(x), value = TRUE)
  mature_testis_cols <- grep("^TS_Mat_", names(x), value = TRUE)
  immature_testis_cols <- grep("^TS_Imm_", names(x), value = TRUE)
  pituitary_cols <- grep("^Pg_", names(x), value = TRUE)
  
  x$Averaged.TPM.ov <- rowMeans(x[, ovary_cols], na.rm = TRUE)
  x$Averaged.TPM.mt <- rowMeans(x[, mature_testis_cols], na.rm = TRUE)
  x$Averaged.TPM.it <- rowMeans(x[, immature_testis_cols], na.rm = TRUE)
  x$Averaged.TPM.pt <- rowMeans(x[, pituitary_cols], na.rm = TRUE)
  
  x <- x[, c("Geneid", "Averaged.TPM.ov", "Averaged.TPM.mt", "Averaged.TPM.it", "Averaged.TPM.pt")]
  
  return(x)
}

###+++###
#Function requires data frame to be normalized
#1. All 0 are set to NA, to exclude them from quatile normalization
#2. Data are quantile normalized
#3. 0 values (the one set to NA) are set back to 0
fQN <- function(x) #
{
	x[x==0] <- NA
	x_m <- as.matrix(x)
	x <- normalize.quantiles(x_m)
	x[is.na(x)] <- 0
	return(data.frame(x))
}	

###+++###	
#Function require a vector with expression of one gene in different tissues.
#Mean is calculated taking in account tissues with 0 expression. 2+0+4=2
fmean <- function(x)
	{
		if(!all(is.na(x)))
	 	{
	 		res <- mean(x, na.rm=TRUE)
	 	} else {
	 		res <- NA
	 	}
	 	return(res)
}

###+++###	
#Function require a vector with expression of one gene in different tissues.
#Max is calculated taking in account tissues with 0 expression. 2+0+4=2
fmax <- function(x)
	{
		if(!all(is.na(x)))
	 	{
	 		res <- max(x, na.rm=TRUE)
	 	} else {
	 		res <- NA
	 	}
	 	return(res)
}

###+++###
#Function require a vector with expression of one gene in different tissues.
#If expression for one tissue is not known, gene specificity for this gene is NA
#Minimum 2 tissues
fTau <- function(x)
{
	if(all(!is.na(x)))
 	{
 		if(min(x, na.rm=TRUE) >= 0)
		{
 			if(max(x)!=0)
 			{
 				x <- (1-(x/max(x)))
 				res <- sum(x, na.rm=TRUE)
 				res <- res/(length(x)-1)
 			} else {
 				res <- 0
 			}
 		} else {
 		res <- NA
 		#print("Expression values have to be positive!")
 		} 
 	} else {
 		res <- NA
 		#print("No data for this gene avalable.")
 	} 
 	return(res)
}

#Calculate adn save tissue specificity Tau#
#1. Data are normalized 
#2. All expression under tpm is set to 0
#3. Replicates mean is calculated (fReplicateMean)
#4. Genes that not expressed in any tissue are removed
#5. Tissue specificity parameters are calculated
tpm <- 1
orgExpression <- na.omit(orgExpression)
print(summary(orgExpression))
nTissues <- length(tissueNames)

##2. if tpm < 1 set as 0
x <- orgExpression[,c(-1)]
x[x < tpm] <- 1
orgExpression[,c(-1)] <- log2(x)
tpm <- log2(tpm)

##3. calculate replicates mean
orgExpression <- fReplicateMean(orgExpression)
orgExpression$Max <- apply(orgExpression[,c(-1)], c(1), fmax)
orgExpression <- orgExpression[orgExpression$Max > tpm,]
orgExpression <- orgExpression[,c(-length(colnames(orgExpression)))]

##4. calculate Tau
orgExpression$Tau <- apply(orgExpression[,c(-1)], c(1), fTau)


##5. add “Preferred_tissue” into table
max_col_index <- max.col(orgExpression[,2:5])
preferred_tissue <- c("ovary", "mature_testis", "immature_testis", "pituitary")[max_col_index]
orgExpression$preferred_tissue <- preferred_tissue

#####-------4. GO enrichment analysis
# prepare GO background 
## input the combined GO backgroud (known genes convert form gProfiler, newly annoatated convert from EggNOG-mapper annotation)
term2gene <- read.table("/ssal_gnpt_rnaseq/analysis/rstudio/term2gene_clean_combine_all_STclassu_emapperGO.txt",sep = " ", header = FALSE)
gene2go = term2gene
names(gene2go) <- NULL

term2name <- read.table("/ssal_gnpt_rnaseq/analysis/rstudio/term2name.txt",sep = "\t", header = FALSE)

termNames = term2name[,c(1,3)]
names(termNames) <- NULL

goterms <- Term(GOTERM)

#convert into a data frame
term2name <- data.frame("GOID"=names(goterms),"term"=goterms )

# add DEG+tau+GM data
degtaugm <- read.table("/ssal_gnpt_rnaseq/analysis/rstudio/DEgenes_tau_WGCNAmodules_integrated.txt", sep = "\t", header = TRUE)
dim(degtaugm)
head(degtaugm)
table(degtaugm$colors) #check WGNCA modules 
table(degtaugm$preferred_tissue) #check perferentialy expressed genes 
table(degtaugm$up_Single) #check single upregulated genes
summary(degtaugm$Tau) #check tau


# enrichments

### turquoise module 
tur <- degtaugm %>%
  filter(colors == "turquoise" & 
           gene_biotype %in% c("novel_protein_coding", "protein_coding"))

dim(tur)

table(tur$gene_biotype)

tur_e = enricher(tur$ref_gene_id,
             gson = NULL,
             TERM2GENE = gene2go,
             TERM2NAME = term2name,
             pAdjustMethod = "BH",
             pvalueCutoff = 0.05)
tur_e

### blue module
blu <- degtaugm %>%
  filter(colors == "blue" & 
           gene_biotype %in% c("novel_protein_coding", "protein_coding"))

dim(blu)

table(blu$gene_biotype)

blu_e = enricher(blu$ref_gene_id,
             TERM2GENE = gene2go,
             TERM2NAME = term2name,
             pAdjustMethod = "BH",
             pvalueCutoff = 0.05)
blu_e

### green module
gre <- degtaugm %>%
  filter(colors == "green" &
           gene_biotype %in% c("novel_protein_coding", "protein_coding"))

dim(gre)

table(gre$gene_biotype)

gre_e = enricher(gre$ref_gene_id,
             gson = NULL,
             TERM2GENE = gene2go,
             TERM2NAME = term2name,
             pAdjustMethod = "BH",
             pvalueCutoff = 0.05)
gre_e

### brown module
bro <- degtaugm %>%
  filter(colors == "brown" &
           gene_biotype %in% c("novel_protein_coding", "protein_coding"))

dim(bro)

table(bro$gene_biotype)

bro_e = enricher(bro$ref_gene_id,
             gson = NULL,
             TERM2GENE = gene2go,
             TERM2NAME = term2name,
             pAdjustMethod = "BH",
             pvalueCutoff = 0.05)
bro_e

### yellow module
yel <- degtaugm %>%
  filter(colors == "yellow" &
           gene_biotype %in% c("novel_protein_coding", "protein_coding"))

dim(yel)

table(yel$gene_biotype)

yel_e = enricher(yel$ref_gene_id,
             gson = NULL,
             TERM2GENE = gene2go,
             TERM2NAME = term2name,
             pAdjustMethod = "BH",
             pvalueCutoff = 0.05)
yel_e

### red module
red <- degtaugm %>%
  filter(colors == "red" &
           gene_biotype %in% c("novel_protein_coding", "protein_coding"))

dim(red)

table(red$gene_biotype)

red_e = enricher(red$ref_gene_id,
             gson = NULL,
             TERM2GENE = gene2go,
             TERM2NAME = term2name,
             pAdjustMethod = "BH",
             pvalueCutoff = 0.05)
red_e

### grey module
grey <- degtaugm %>%
  filter(colors == "grey" &
           gene_biotype %in% c("novel_protein_coding", "protein_coding"))

dim(grey)

table(grey$gene_biotype)

grey_e = enricher(grey$ref_gene_id,
             gson = NULL,
             TERM2GENE = gene2go,
             TERM2NAME = term2name,
             pAdjustMethod = "BH",
             pvalueCutoff = 0.05)
grey_e

### pink module
pink <- degtaugm %>%
  filter(colors == "pink" &
           gene_biotype %in% c("novel_protein_coding", "protein_coding"))

dim(pink)

table(pink$gene_biotype)

pink_e = enricher(pink$ref_gene_id,
             gson = NULL,
             TERM2GENE = gene2go,
             TERM2NAME = term2name,
             pAdjustMethod = "BH",
             pvalueCutoff = 0.05)

### black module
black <- degtaugm %>%
  filter(colors == "black" &
           gene_biotype %in% c("novel_protein_coding", "protein_coding"))

dim(black)

table(black$gene_biotype)

# enrichment
black_e = enricher(black$ref_gene_id,
             gson = NULL,
             TERM2GENE = gene2go,
             TERM2NAME = term2name,
             pAdjustMethod = "BH",
             pvalueCutoff = 0.05)
black_e

##! there were six moudles had enriched terms
# extrct newly annoatated genes enriched in different modules
tur_e_novelgene <- tur_e[grepl("MSTRG.", tur_e$geneID), ]
blu_e_novelgene <- blu_e[grepl("MSTRG.", blu_e$geneID), ]
bro_e_novelgene <- bro_e[grepl("MSTRG.", bro_e$geneID), ]
yel_e_novelgene <- yel_e[grepl("MSTRG.", yel_e$geneID), ]
gre_e_novelgene <- gre_e[grepl("MSTRG.", gre_e$geneID), ]
black_e_novelgene <- black_e[grepl("MSTRG.", black_e$geneID), ]
```
